# The Use of GC-, Codon-, and Amino Acid-frequencies to Understand the Evolutionary Forces at a Genomic Scale

**DOI:** 10.1101/863142

**Authors:** Arne Elofsson

## Abstract

It is well known that the GC content varies enormously between organisms; this is believed to be caused by a combination of mutational preferences and selective pressure. Within coding regions, the variation of GC is more substantial in position three and smaller in position one and two. Less well known is that this variation also has an enormous impact on the frequency of amino acids as their codons vary in GC content. For instance, the fraction of alanines in different proteomes varies from 1.1% to 16.5%. In general, the frequency of different amino acids correlates strongly with the number of codons, the GC content of these codons and the genomic GC contents. However, there are clear and systematic deviations from the expected frequencies. Some amino acids are more frequent than expected by chance, while others are less frequent. A plausible model to explain this is that there exist two different selective forces acting on the genes; First, there exists a force acting to maintain the overall GC level and secondly there exists a selective force acting on the amino acid level. Here, we use the divergence in amino acid frequency from what is expected by the GC content to analyze the selective pressure acting on codon frequencies in the three kingdoms of life. We find four major selective forces; First, the frequency of serine is lower than expected in all genomes, but most in prokaryotes. Secondly, there exist a selective pressure acting to balance positively and negatively charged amino acids, which results in a reduction of arginine and negatively charged amino acids. This results in a reduction of arginine and all the negatively charged amino acids. Thirdly, the frequency of the hydrophobic residues encoded by a T in the second codon position does not change with GC. Their frequency is lower in eukaryotes than in prokaryotes. Finally, some amino acids with unique properties, such as proline glycine and proline, are limited in their frequency variation.

## 2 Introduction

The GC-frequency varies significantly between different and within genomes, both in coding and non-coding regions [1]. The reason behind this is not entirely understood. However, it is likely due to a combination of a balance between mutational preferences, selective pressure and evolutionary history [1]. In general mutational preferences decrease GC levels in most organisms [2, 3], while GC-biased gene conversion (gBGC) can contribute to higher GC levels [4]. The environment of the organism can also influence the GC level as GC levels are higher in thermophiles [5]. Further, there is a phylogenetic signal so that closely related organisms mostly have similar GC-levels [6]. Finally, differences in DNA polymerase subunit III might correlate with differences in GC [7].

Here, we are not trying to answer the long-disputed origin of the difference in GC content. Instead, we assume that there is some mechanism driving the GC content of a particular organism towards an optimal level. Thereafter, we ask how does this affect the proteomes by examining the frequency of amino acids, nucleotides and codons. A different number of codons encodes the different amino acids. These codons differ in GC content. Therefore, in general, amino acids encoded by more codons are more frequent, and amino acids encoded by GC-rich codons are more frequent in GC-rich genomes [8–10] leading to massive variation in the frequency of amino acids in different organisms [11, 12]. For instance, the positively charged amino acids Arg and Lys vary between 2% and 10% in frequency. Arg is more frequent in GC-rich organism, and Lys is more frequent in GC-poor organisms.

The codon table, see Figure 1, is surprisingly well conserved since early life. The same 61 codons encode the twenty amino acids in most organisms. However, some variations exist. For instance, in eukaryotes, one of the stop codons can encode selenium methionine [13], and other variations exist among Mycoplasma, Spiroplasma, Ureaplasma and Mesoplasma [14]. The redundancy in the codon table means that for many amino acids, the third position does not change the amino acid. Therefore, the overall GC content can change by using different nucleotides in the third position without affecting the proteome. Further, the codons have evolved in such a way that the general properties of the amino acids are determined mainly by the codon in position two [15].

**Figure 1.**
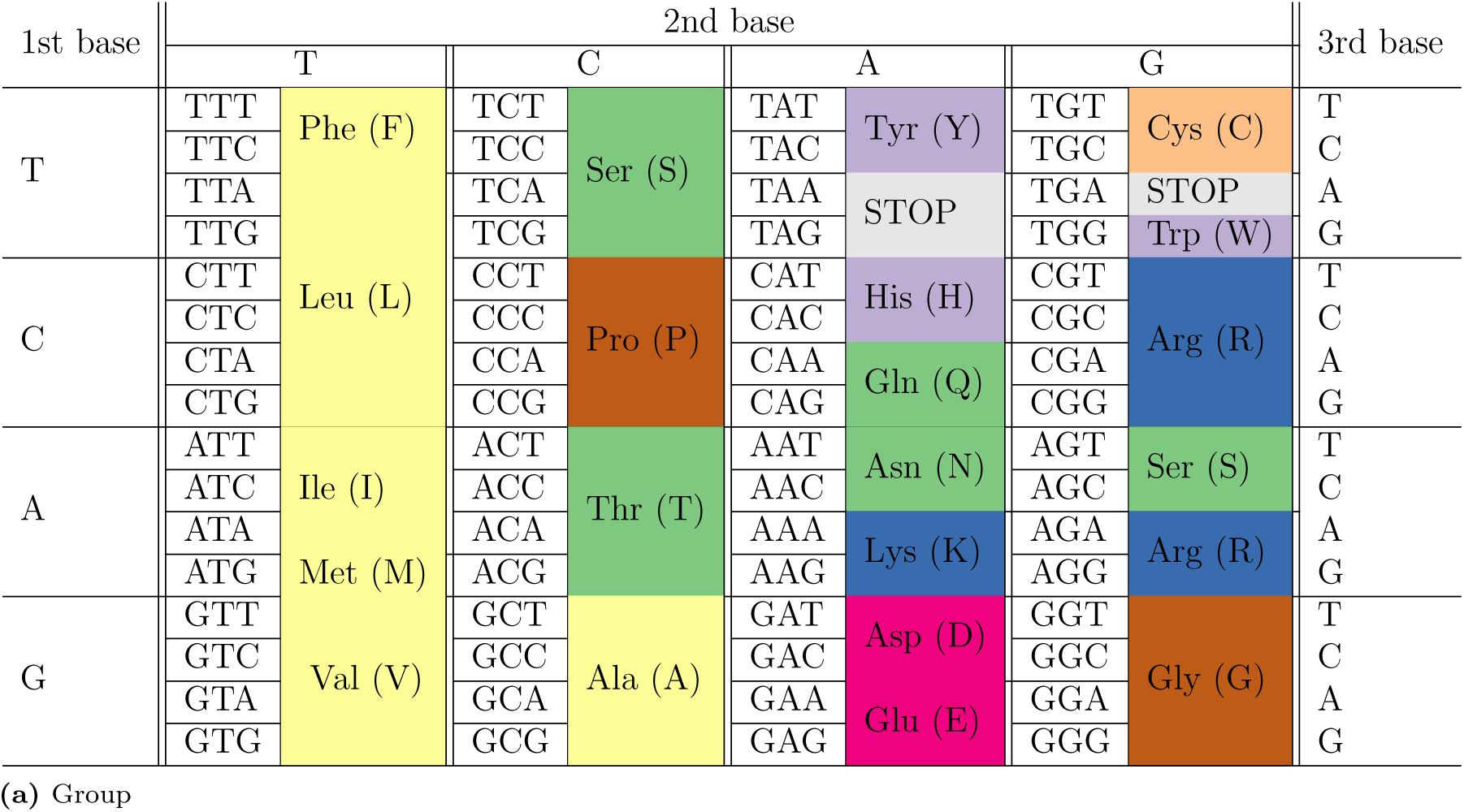
Codon tables with the amino acids encoded according to different properties. (a) The colour is based on the amino acid type (hydrophobic - yellow, Basic - blue, Acidic - red, Polar - green, amphipathic - purple and loop-preferring brown) (b) coloured according to pI-values to be neutral (c) coloured by secondary structure preference and (d) coloured according to disorder preference. The figure is inspired by a figure at Wikipedia at http://www.wikipedia.org/

In addition to codon frequency and GC level, there exist many factors that contribute to the frequency of amino acids [16–18]. The cost of amino acid synthesis might affect their frequency [19, 20], and some amino acids, such as serine, can be toxic at high levels [21]. In addition to purifying effects to reduce the frequency of one amino acid, an organism might require a minimum frequency of amino acids with specific properties, while other amino acids, such as alanine, might be allowed to vary more freely [22].

Below, we analyze the frequency of amino acids, codons and nucleotides in different genomes. We show that the codon frequency does not fully explain amino acid frequencies, i.e. other factors also affect the amino acid frequencies. Some amino acids, such as serine, are consistently less frequent than expected, while others, such as glutamate, are more frequent. Further, some amino acids, such as proline, are less dependent on GC than expected, indicating that there are limits to how much they can vary. The picture that emerges is that there on a genomic perspective there exists two selective forces, one that adjusts the GC content to a certain level and one that given a certain GC level adjusts amino acids frequencies. By detailed analysis, we can obtain an understanding of the forces acting on the amino acids.

## 3 Material and Methods

### 3.1 Datasets

The dataset used in this study originates from the complete bacterial, archaeal and eukaryotic proteomes in UniProt [23] as of December 2017. All genomes from Mycoplasma, Spiroplasma, Ureaplasma, and Mesoplasma were ignored as they have another codon usage - which influences the expected amino acid frequencies. The final dataset contains 36,098,162 protein sequences from 8,546 genomes, divided into 7,197 bacterial, 351 archaeal, and 998 eukaryotic species. For each genome, the GC content of the genome and the length was obtained from NCBI. Further, the DNA and amino acid sequences of each gene were downloaded. The processed datasets, as well as all scripts, are available from this repository [24].

### 3.2 Statistics

For each protein, we calculated amino acid-, GC-, codon- and nucleotide-frequencies. Average, maximum and minimum frequencies for each genome in the dataset are presented in Table S1.

ANOVA type 2 F-tests [25] were used to identify the contribution differences between the kingdoms, compensating for differences in GC content, see Table **??**. Using each codon/amino acid/nucleotide as the dependent variable and the GC content is used as the independent variable, the difference between kingdoms was tested. Here, it should be mentioned that even tiny differences are statistically significant, as the dataset is large. Further, differences between eukaryotes and bacteria dominate the ANOVA test as these are the most prominent groups.

### 3.3 Expected frequencies

It is necessary to define the expected frequency of amino acid (*AA*^*i*^) to identify any selective pressure. Therefore, we define models to estimate the expected amino acid frequencies 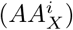 assuming different scenarios. The simplest model, the *codon* model, assumes that the frequency of an amino acid is solely determined by the number of codons encoding that amino acid:

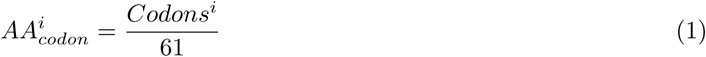

where *Codons*^*i*^ is the number of codons for amino acid *i* and 61 is the number of codons excluding stop codons.

Alternatively, the amino acid frequencies may be dependent on GC (i.e. there exist some other mechanism that determines the GC content of a genome) leading to the expected amino acid frequency 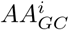 at a certain GC level to be:

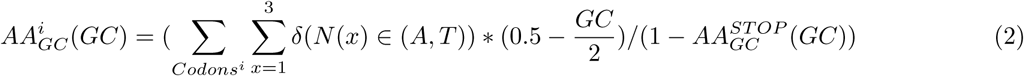

where *Codons*^*i*^ represents the codons for amino acid *i* and *x* the three nucleotides in that codon, GC is the fraction GC in the genome and *δ*(*N*(*x*) ∈ (*A, T*)) is a delta function that is one if the nucleotide *N*(*x*) is A or T, and zero if not. Further, 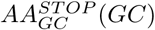 is the expected frequency of stop-codons given GC as defined here:

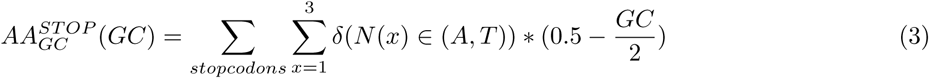

where the ∑_*stopcodons*_ sums over the three stop codons.

However, as we show below there are other parameters that also affect the amino acid frequencies. To take several scenarios into account, we use the following formulae to estimate the amino acid frequency (*AA*^*i*^) for the amino acid *i* at a given *GC* level:

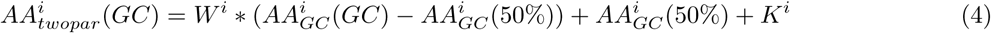

Here, 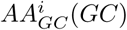 is the expected frequency of amino acid *i* at the GC as defined in equation 2. 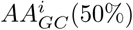 is the expected frequency at GC=50%, and *W*^*i*^, and *K*^*i*^ are two parameters that are optimized for each amino acid. The reason to use this function, and not simply *AA*^*i*^ = *w*^*i*^ * *GC* + *k*^*i*^ is to have a consistent definition of the parameters *W*^*i*^, and *K*^*i*^. In particular, the parameter *K*^*i*^ is useful to estimate over-, and under-representation of an amino acid.

Using equation 4, we can model different scenarios. If *W*^*i*^ = 0 and *K*^*i*^ = 0, then equation 4 describes the expected frequency from the number of codons as in equation 1 (the *codon* model). If *W*^*i*^ = 1 and *K*^*i*^ = 0 the equation describes the expected amino acid frequency at a certain GC level as in equation 2. If *W*^*i*^ = 0 while *K*^*i*^ is optimized, this describes the average amino acid frequency in all genomes and then *K*^*i*^ represents a shift from the expected frequency. Finally, we can optimize both *W*^*i*^ and *K*^*i*^ and obtain the amino acid levels using two parameters (the *twopar* model). Here, to be more realistic, we limit *W*^*i*^ to be between 0 and 1. Also here *K*^*i*^ represents the shift from the expected frequency.

To compare the different models to estimate the amino acids, we use the Pearson correlation coefficient [26] and the average error between the estimated and observed frequencies of all twenty amino acids, see Figure S1.

### 3.4 Linear regressions

To estimate the GC frequency from amino acid frequency, we used sklearn [27]. Given the amino acid frequency of one or more amino acids in a protein or a proteome, the model was trained to predict the GC level of the coding region of a proteome. In addition we trained the same model to predict the GC level from a single protein. Here 25,000 randomly selected proteins were used.

## 4 Results and Discussion

The GC frequency can vary tremendously between organisms, see Figure 2. In our set of proteomes, the beta proteobacteria *Candidatus Zinderia insecticola* has the lowest GC content with 13.5% and *Geodermatophilus nigrescens* has the highest (75.9%), see Table S1. The mechanism causing these differences is not entirely known, but factors such as mutation rate, crossover rate, thermodynamical stability and phylogenetic memory contribute [4]. Anyhow, in this study, we will not focus on the GC difference. Instead, we will analyze how the difference in GC levels affect the proteomes and use divergence from expected frequencies to analyze the selective pressures at the proteome level.

**Figure 2.**
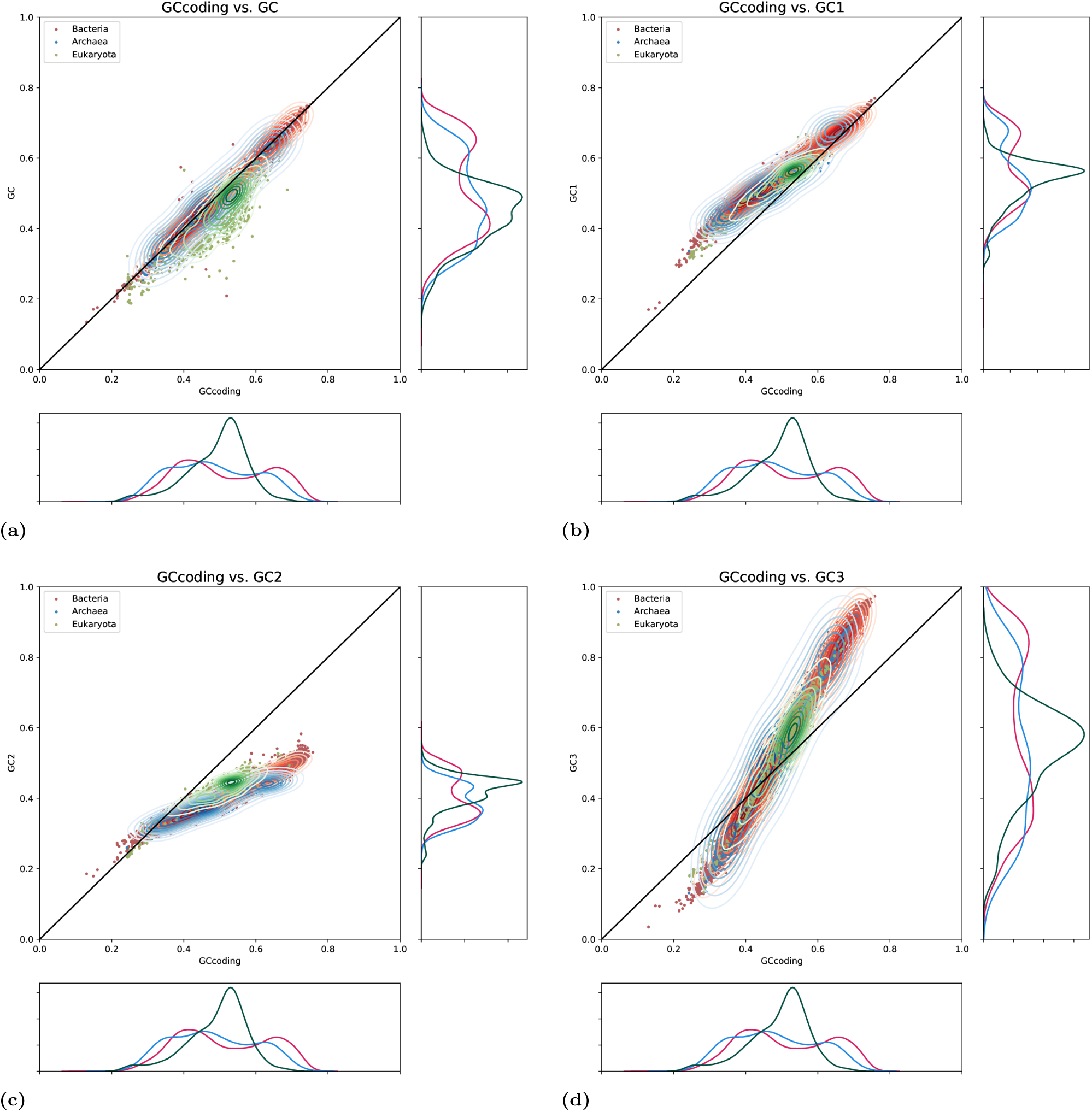
Distribution of GC contents in the three different kingdoms. In (a) the GC content in the whole genome is plotted against the GC content in the coding regions. In (b-d). the GC content in the three codon positions are plotted against the GC in the coding regions.

### 4.1 GC distributions

First, some notes about the overall GC content. Both prokaryotic kingdoms have a bimodal GC distribution with one peak around 40% and the second at 70% [28], see Figure 2a. In contrast, Eukaryotes have a single, less wide, peak of GC content. Therefore, the standard deviation in the prokaryotes is larger (12% vs 8%), while the average GC levels are similar (48%-51% for the coding regions), see Table S1.

#### 4.1.1 GC coding vs non-coding

For both prokaryotes, the genomic GC level (from NCBI) and the GC level of the coding regions (from Uniprot) are almost identical and perfectly correlated (CC=0.998), while for Eukaryotes the levels differ slightly but are still strongly correlated (CC=0.89), see Table S1. Eukaryotes have a higher GC content in coding regions (49% vs 44%), see Figure 2a. The GC level in the coding regions is similar to the average level observed in prokaryotes. Therefore, we believe that comparing features with the GC content of the coding region is more appropriate. Further is also simplifies the analysis of codon and nucleotide frequencies.

### 4.2 The selective pressure at the GC level

In the codon table, seven (Phe, Leu, Val, Pro, Thr, Ala, and Gly) out of the twenty amino acids are determined by position one and two, see Figure 1. Further, the two first bases and a combination of TC or AG in position three determines eight other amino acids (Tyr, His, Gln, Asn, Lys, Asp, Glu, and Cys). Two amino acids (Met and Trp) have only one codon, and Ile uses the three ATX codons not encoding Met. The remaining two amino acids, serine and arginine, are encoded by two groups of codons with different nucleotides in position one and two. Finally, there are three stop codons that all have a T in its first position (T1).

Given the position in the codon table for amino acids with similar properties, it is clear that in particular, position two determines the properties of the amino acid [15]. For instance, all codons with T2 encode hydrophobic amino acids, while both negatively charged amino acids have A2.

#### 4.2.1 GC in different positions

The GC content differs between the three codon positions, see Figure 2 and Table S1. In all positions, the GC content is strongly correlated with the overall GC content (*Cc* > 0.93). The average GC content is lower in position two than in the other two positions. Further, in position one and two, the variation in GC content is much more restricted than in position three. The highest GC level in position three is 97% and the lowest 3%, compared to 18% and 58% in position two. The difference between the positions means that the GC variation in position three is significantly higher than in the other parts of the genome.

A model to explain the variation of GC in the three positions can be formulated as follows: In an organism, there exists a selective pressure to have a certain optimal GC content (in the coding regions). For some organisms, this optimal level is very high or very low, i.e. extreme. However, the selective pressure acting on amino acids frequencies makes it impossible to have extreme GC levels in position one and two. Therefore, to obtain extreme overall GC levels in these organisms, it is necessary to over-compensate in position three. In theory, if the GC content is limited to 50% in position one and two but varying in position three, this allows the genomic GC to vary between 33 and 67%. However, the GC content in one and two also varies, and amino acid frequencies also change; therefore, the overall GC content can vary between 13% and 76%.

Although the average GC content is similar in all three positions, it is clear that the nucleotide frequencies are not, see Figure 3, 4 and Table S1. The differences are largest for position one and two. In position one, G1 and A1 are more frequent than the other two nucleotides, while in position two A2 and T2 are most frequent. For a more detailed understanding of these differences, we will analyze the frequencies of each nucleotide in each position, starting with position one.

**Figure 3.**
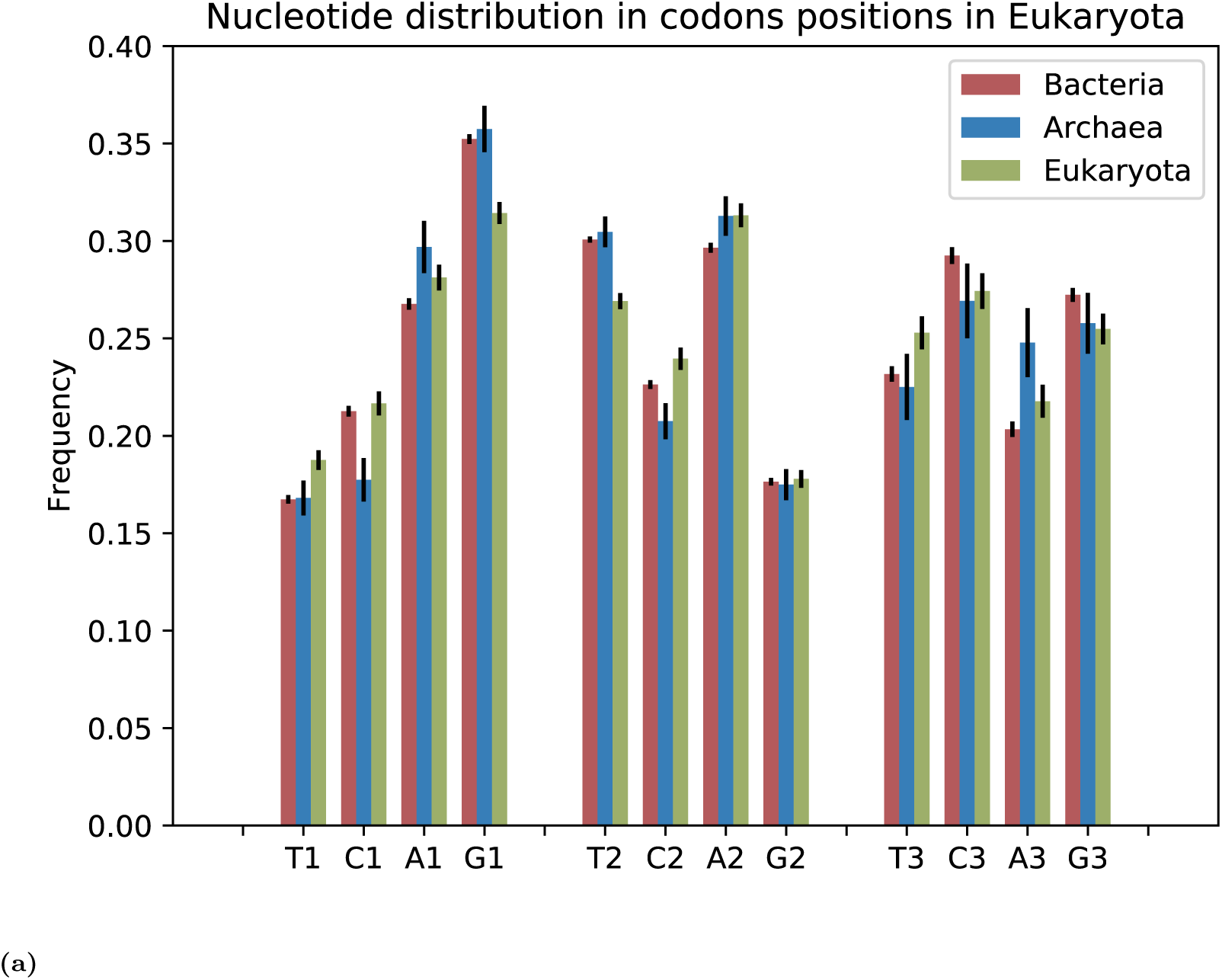
Average composition of nucleotides in different codon positions.

**Figure 4.**
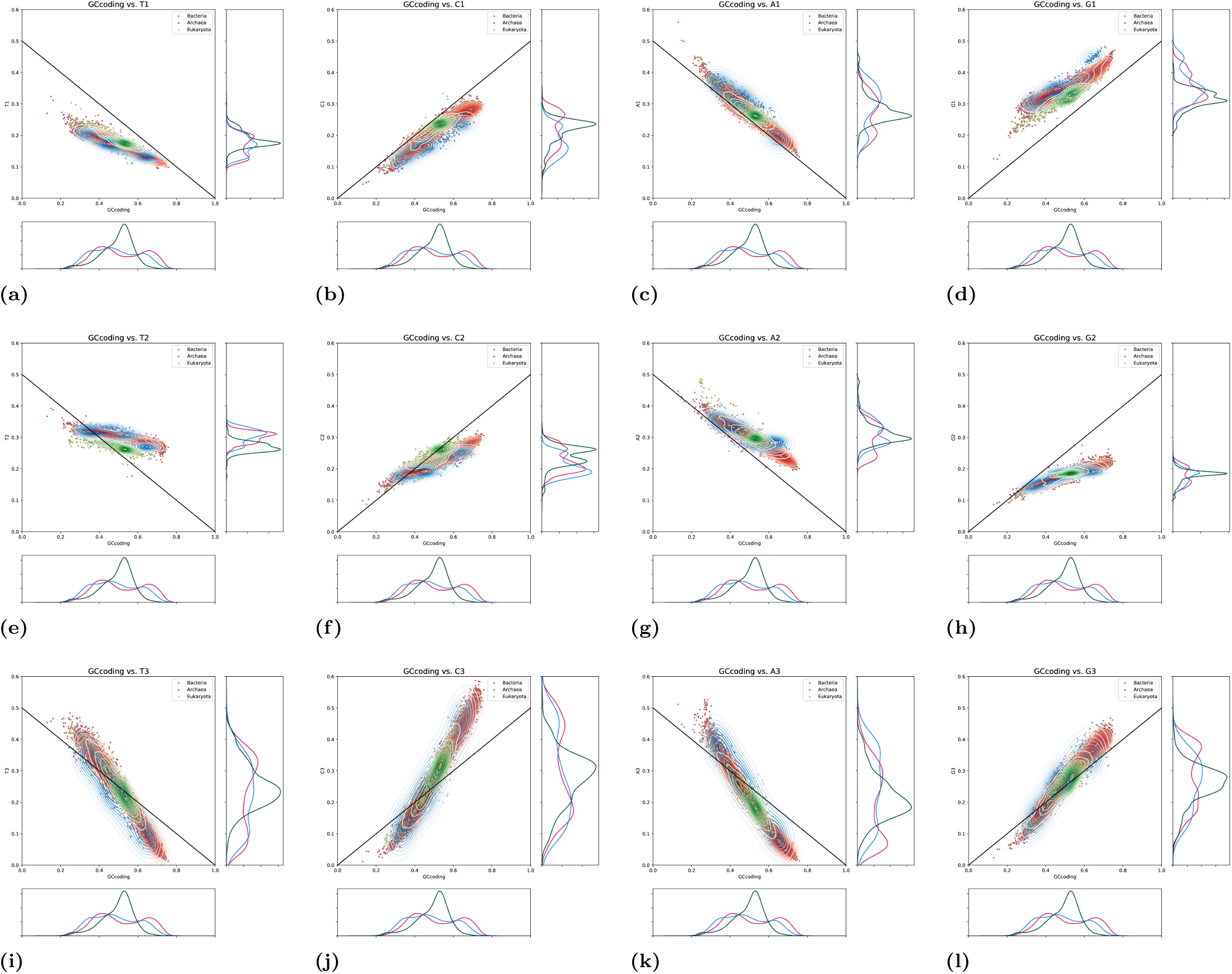
Position specific nucleotide frequencies plotted against the GC frequency.

#### 4.2.2 Position 1

In position one, it can be seen that G1 is most frequent, and T1 is least frequent (average frequency is 16.7%). However, it should be remembered that all three stop codons have a T1, so the expected T1 frequency is not 25% but only 20.3%. In addition, serine, which has four out of six codons with T1, is one of the most underrepresented amino acids, as we have described before [12]. G1 encodes for VADEG, these amino acids are all over-represented compared to random, see Figure 5.

**Figure 5.**
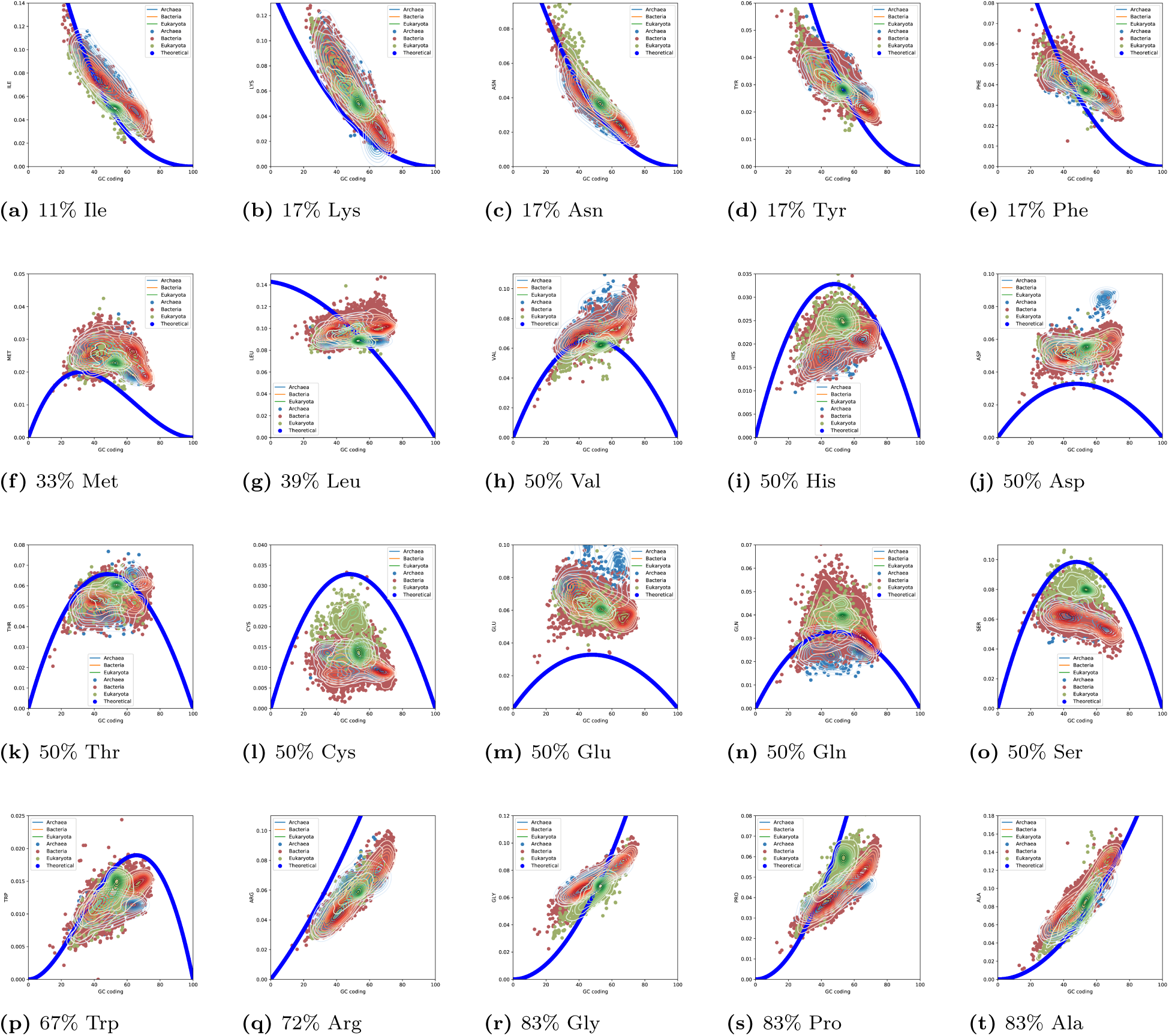
Frequency of different vs GC of the genomes amino acids are sorted by the GC content of the codons. The amino acids are sorted by their TOP-IDP scores. The number represent the fraction of GC among the codons. The blue line represent the expected fraction according to the codon frequency. The purple lines represent the expected fraction from the codon position GC content.

#### 4.2.3 Position 2

Position two is the most conserved position when it comes to GC content. It is also clear that A2 and T2 are more frequent than G2 and C2, on average about 30% vs 20%, see Table S1 and Figure 3. Further, T2 is almost independent of the GC content in all genomes, but consistently lower in eukaryotes than prokaryotes, see Figure 4. The constant level of T2 guarantees a stable amount of the non-polar, and *β*-sheet forming amino acids (FLIVM). G2 is rare and have a limited range. G2 encodes several amino acids with unique properties, such as glycine (the smallest amino acid) and cysteine (that can form disulphide bonds), but also arginine, tryptophan and one of the stop codons. The rareness can be contributed to the 40% (6 out of 15) of the non-stop G2 codons that encode arginine, and the frequency of arginine is underrepresented, see Figure 5. Finally, A2, that encodes primarily charged and polar amino acids (YHQNEDK), and C2, that encodes APST are allowed to vary more freely than the other two nucleotides in position 2.

#### 4.2.4 Position 3

In general, it is believed that position three in a codon is not under selective pressure as it only rarely affects the amino acid, Figure 1. However, if no selective pressure acted on position three random drift would make all nucleotides equally frequent in that position, and clearly, they are not, see Figure 4. In contrast, the nucleotides in position three vary much more than in the other positions. The frequency of most nucleotides varies between 1% and 60%, supporting the idea that the genomic GC preference governs nucleotide frequencies. C3 is most frequent in position three but least frequent in the other two positions, see Figure 3.

### 4.3 Amino acid frequency vs GC

To be able to identify the selective pressures acting on amino acid frequencies, it is necessary to estimate the expected amino acid frequencies without any selective pressure. Therefore, it is necessary to have a model to describe the expected amino acid frequency for a genome. Following the speculations above we do assume that: There exist some evolutionary process that strives the GC content of a genome to be adapted, but that is independent of the selective pressure acting at the amino acid level. It is then possible to model the expected amino acid frequencies assuming that protein-coding regions would be random in the absence of any selective pressure at the amino acid level.

A simple explanation of the variation of amino acid frequency would be that it is just decided by the number of codons coding for an amino acid as in equation 1. Nevertheless, a better agreement is observed when taking the GC into account and calculate the expected amino acid frequencies, given the GC of the genome, as in equation 2. Below, we use this equation to estimate the expected frequencies of the amino acids.

Figure 5 shows the amino acid frequencies of each amino acid against the GC content of the coding regions with the blue lines representing the expected amino acid frequencies according to equation 2. The sorting of the amino acids is based on the GC content in their codons.

#### 4.3.1 Frequency of low GC amino acids depends strongly on GC

The frequency of all the amino acids with less than one-third of GC in their codons, i.e. Ile, Lys, Asn, Phe and Tyr, show a strong correlation with GC, see the top row in Figure 5. The frequency of these amino acids vary from 1-2% at high GC up to 19% at low GC and the correlation with GC is 0.83 to 0.93, see Table S1. The lowest correlations against GC are for Tyr and Phe, which have a flatter distribution than expected from the GC frequency alone.

#### 4.3.2 The frequencies of amino acids low GC dependency are independent of GC

Next, there are 11 amino acids with a GC content in their codons between one- and two-thirds. None of these shows a strong dependency of GC, but the correlations with GC are rather high for Valine (CC=0.72) and Trp (CC=0.74). More notably, some of these amino acids are more frequent than expected from the codons and some less.

#### 4.3.3 Frequency of all high GC codons strongly depends on GC

Finally, the amino acids with more than two-third of GC in their codons are also strongly dependent on the GC content (CC*>* 0.85). Shifts can be seen as Arg is less frequent than expected. The frequencies of Gly and Pro also appears to be limited to be within a specific range.

### 4.4 Systematic shifts

From the studies above, it is clear that there exist systematic divergences of amino acid frequencies for some amino acids. In general, the divergences are (a combination) of two types, shifts and decreased GC dependency. A shift refers to that the amino acid frequency is consistently over- or under-represented (as for serine), while the decreased GC dependency refers to a decreased dependency of GC, i.e. a flatter distribution (as for proline), see Figure 5. It is tempting to speculate that a shift would indicate that there exist a selective pressure for that amino acid to be more or less frequent, while a decreased GC dependency indicates that there exists a selective pressure to keep that amino acid at a constant level.

To identify systematic shifts, we have used equation 4 (with different limitations to the parameters). Here, *K* describes a shift up or down from what is expected by random and *W* describes the strength of the dependency with GC (one is perfectly correlated, and zero indicates no dependency). The parameter *W* is, therefore, only relevant for the amino acids with GC-rich or GC-poor codons, see Figure S3.

Figure 6 shows that the shifts (*K*) are consistent independent of what model is used. Arginine, serine, cysteine and proline are under-represented while glutamate, aspartate, lysine and alanine are over-represented, see Figure 6. These shifts are also clearly observable in Figure 5. The variation between the kingdoms is small, but the shifts are in general smaller in Eukaryotes. The average error for the GC model, equation 2, is 1.4% in Eukaryotes vs 1.9% in Bacteria and 2.1% in Archaea.

**Figure 6.**
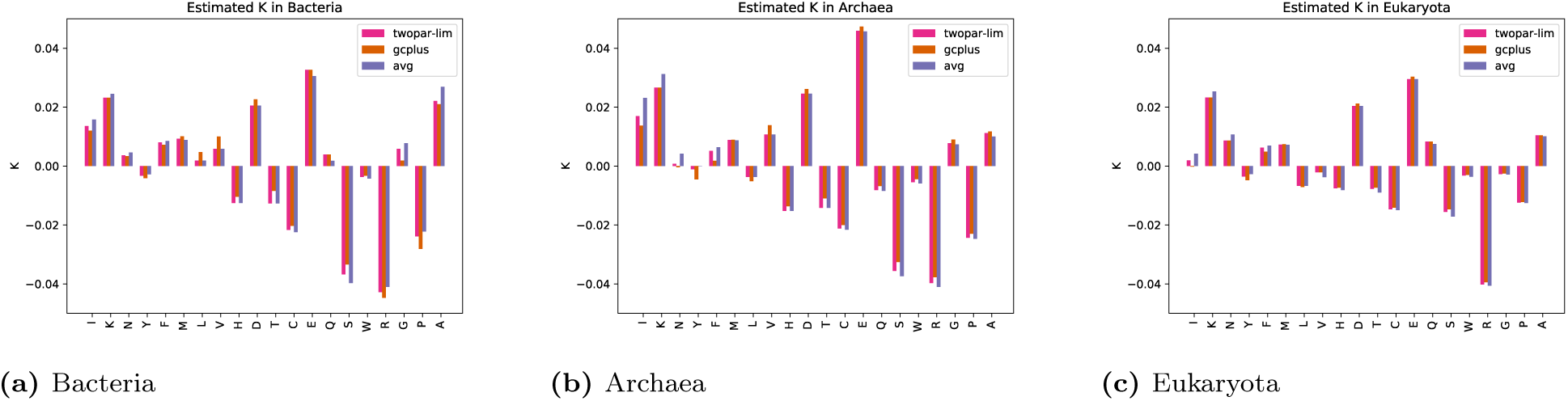
Combined frequency plots

### 4.5 The intricate balance of charged residues

The positively charged amino acids Lys and Arg are like Siamese twins, one has GC-rich codons, and one GC-poor, both are positively charged, and they can often (but not always) perform similar roles in a protein. One notable difference is that six codons encode arginine compared with two for lysine, i.e. arginine should be three times as frequent at 50% GC. However, arginine is consistently less frequent than expected from GC while lysine is more frequent, compensating for the difference in codons, see Figure 5 and 6. The total number of Arg+Lys is rather constant but decreases slightly with GC, see Figure 7.

**Figure 7.**
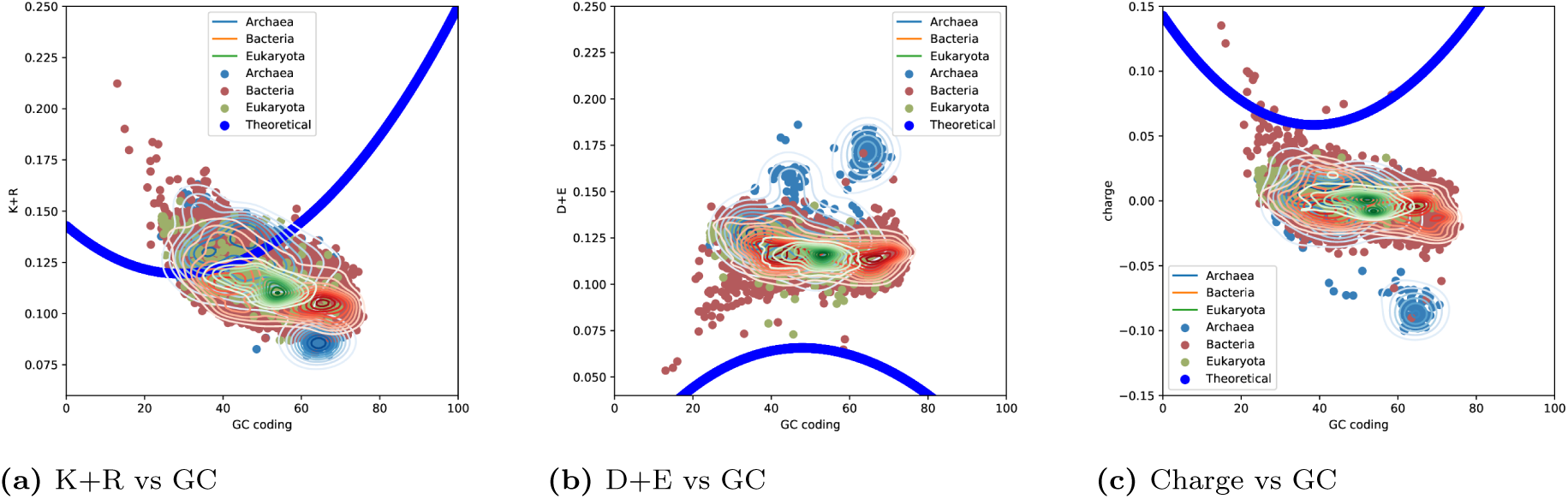
Frequencies of groups of amino acids vs GC.

The negative amino acids (Asp and Glu) are, in contrast, not very GC dependent and are constant in GC, see Figure 5. Notably, as a group, the negatively charged amino acids are much more frequent compared to what is expected by random, see Figure7 and 6. The shifts are therefore most likely a consequence of that there are eight codons for positively charged amino acids compared to only four for the negatively charged amino acids and that the overall charge of the proteome is close to neutral independent on GC content, see Figure 7.

### 4.6 Limited frequency ranges

In addition to amino acids consistently over- or under-represented, there exist amino acids that are limited in their variation. In Figure 5 and S3, it can be seen that five amino acids are less dependent on GC than expected. Isoleucine, tyrosine and phenylalanine are all less frequent than expected at low GC and more frequent at high GC. Similarly, Pro and Gly are both more frequent than expected at low GC and less frequent at high GC. Given the unique properties of Tyr/Phe (aromatic) and Gly/Pro (secondary structure breakers), it is not surprising that there exist boundaries to their frequency variations.

### 4.7 Differences between kingdoms

Although most amino acids and codon frequencies are similar in the three different kingdoms, there exist some differences to be noted. We have earlier reported that serine and proline are more frequent in eukaryotes, and that isoleucine is less frequent [12]. Here, we confirm that these differences are among the most significant differences between the kingdoms using an ANOVA test, see Table 1. However, other differences can also be detected.

**Table 1.**
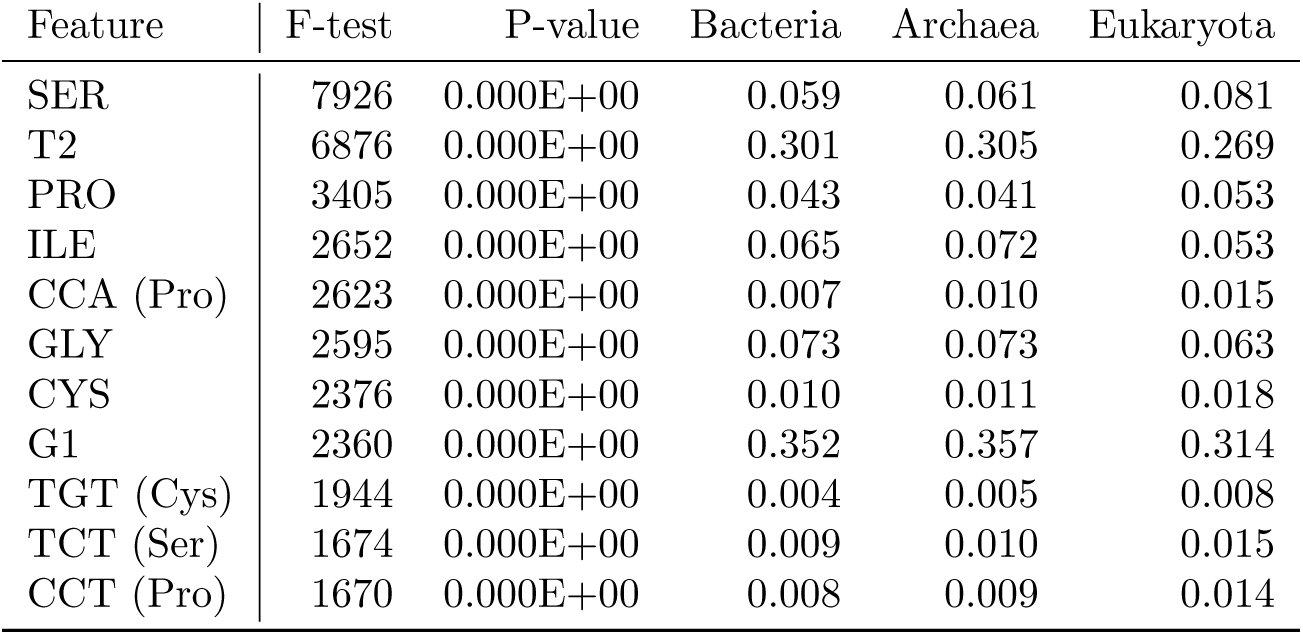
ANOVA tests for comparison between kingdoms. The most significant features when comparing all three kingdoms are listed here, for all other comparisons, see supplementary Table **??**. The average frequencies feature in the three kingdoms are shown. Note that the average feature reported here does not compensate for differences in GC contents as done in the ANOVA test.

In Table 1, it can be seen that two features dominate the difference between the kingdoms, increased serine frequency in eukaryotes and decreased T2 frequency in eukaryotes. As mentioned above, T2 codes for the hydrophobic amino acids phenylalanine, leucine, isoleucine, methionine and valine.

If we ignore differences in codons, following next in importance is the increase in eukaryotic proline frequency and decrease in isoleucine frequency [12]. These are then followed a decrease in glycine and an increase in cysteine in the eukaryotes. Finally, G1 is less frequent in eukaryotes than in prokaryotes. G1 encodes valine, alanine, aspartate, glutamate and glycine, and all these are slightly less frequent in eukaryotes than in the prokaryotes.

All the codons that are highest ranked in the ANOVA test are coding for one of the amino acids discussed above. It is interesting to note that CCA codon explains the most of the proline increase.

#### 4.7.1 Archaea

For many features, such as glutamate and aspartate frequencies, it can be seen that the archaea kingdom is divided into two groups. Brief analysis indicates that this roughly correlates with the phylum Euryarchaeota and other archaea. Euroarchaeota have more proteins (2170 vs 1620), higher GC (50% vs 45%) and more Asp (6.3% vs 4.9%) and Glu (8.2% vs 7.2%) but less Lys (7.3% vs 5.8%). Although interesting, a detailed analysis of these differences is beyond the goals of this study.

### 4.8 Predicting GC from amino acid frequencies

Is it possible to predict the GC frequency from amino acid frequencies? We show that even the frequency of one amino acids, such as asparagine or alanine, in the proteome, can predict the GC level with an error of less than 5% and a correlation coefficient of 0.95, see Figure 8. If the frequency of all twenty amino acids is included, the error drops below 2%, and the correlation coefficient is 0.99.

**Figure 8.**
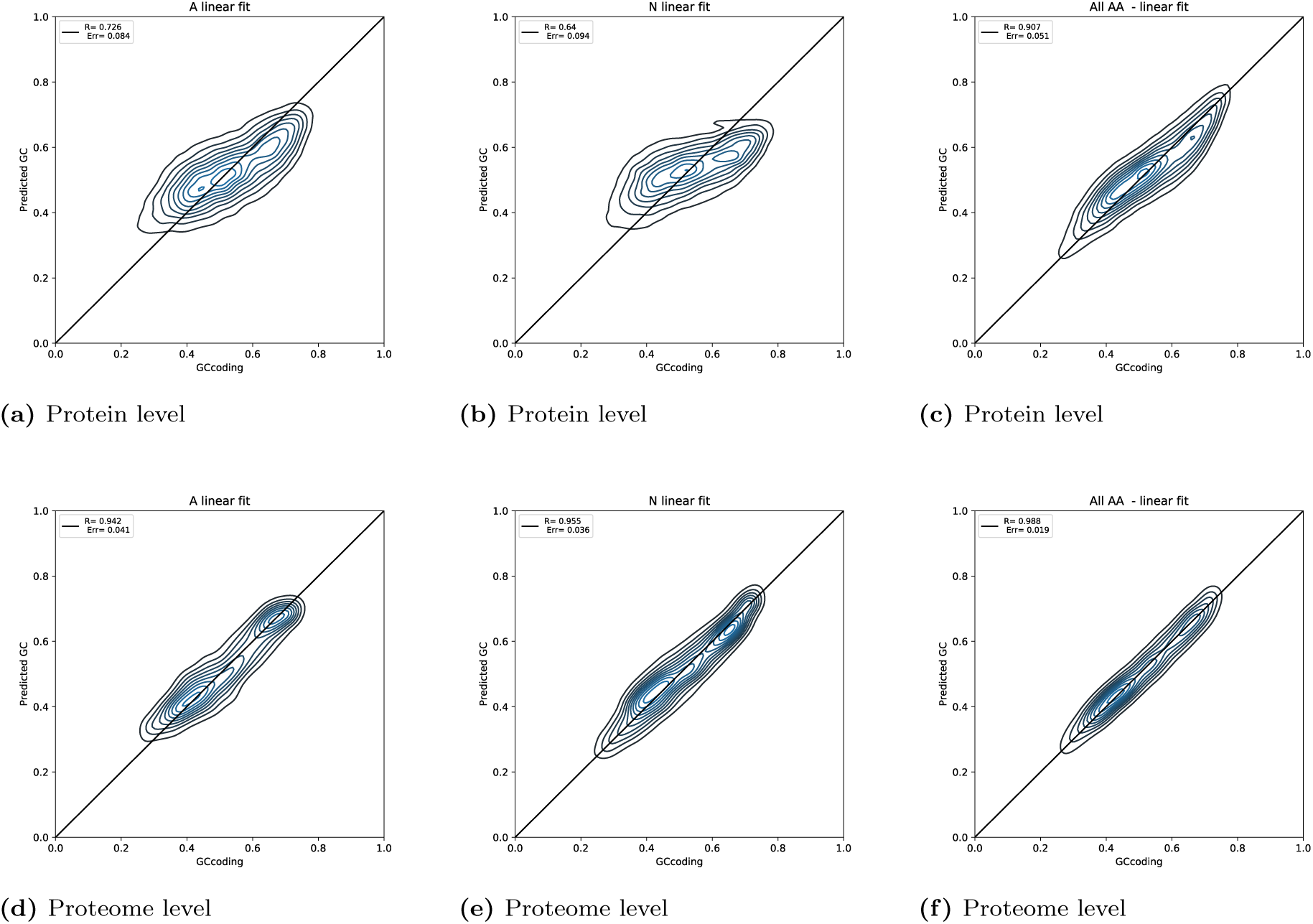
Predicting GC content from amino avid frequencies.

Even the frequency of amino acids for a single protein is informative of the GC level of the entire proteome. The sequence of a single protein can predict the GC level with an average error of 5% and a correlation coefficient above 0.90. This can, for instance, be used to detect laterally transferred genes directly from amino acid sequences if the genomic sequence was not available.

## 5 Conclusions

Here, we study the relationship between GC content of organisms and frequencies in their coding regions. We highlight that amino acid frequencies differ significantly in high and low-GC genomes and that their frequencies are primarily dependent on the number of codons and the GC content of their codons. But there are also significant differences.

To explain this, we propose that there exist an (unknown) mechanism acting to maintain the GC level in an organism. This can be seen by the fact that the third position varies much more than the others and by the differences in nucleotide frequencies in the different codon positions. Next, we propose that there is also a selective pressure changing amino acid frequencies from what is expected by chance. This mechanism decreases the frequencies of arginine and serine in all organisms, while lysine, aspartate and glutamate are more frequent than expected by chance. Further, this mechanism limits the influence of GC on the frequency of tyrosine, phenylalanine, glycine, proline and isoleucine.

We also note that the selective pressure acts to; (i) Keep a balance of negatively and positively charges amino acids in all genomes (except some Euroarchaeota). This is maintained by an intriguing by the underrepresentation of arginine and overrepresentation of negatively charged amino acids. (ii) Maintaining the hydrophobic residues at a constant level by keeping a constant fraction of Thymine in the second codon position.

Finally, we also show that two most significant factors differ between eukaryotes and prokaryotes are: (a) Eukaryotes have more serine residues and (b) less of codons with a T in position two (T2), which results in fewer hydrophobic residues (FLIVM).

## Acknowledgements

We do thank Lukas Käll, Lars Arvestad, Jens Lagergren and Patrick Bryant for valuable inputs and discussions. We do also thank for valuable discussions with the members of the COST Action BM1405 NGP-net.

## 6 Supplementary Material

**Figure S1.**
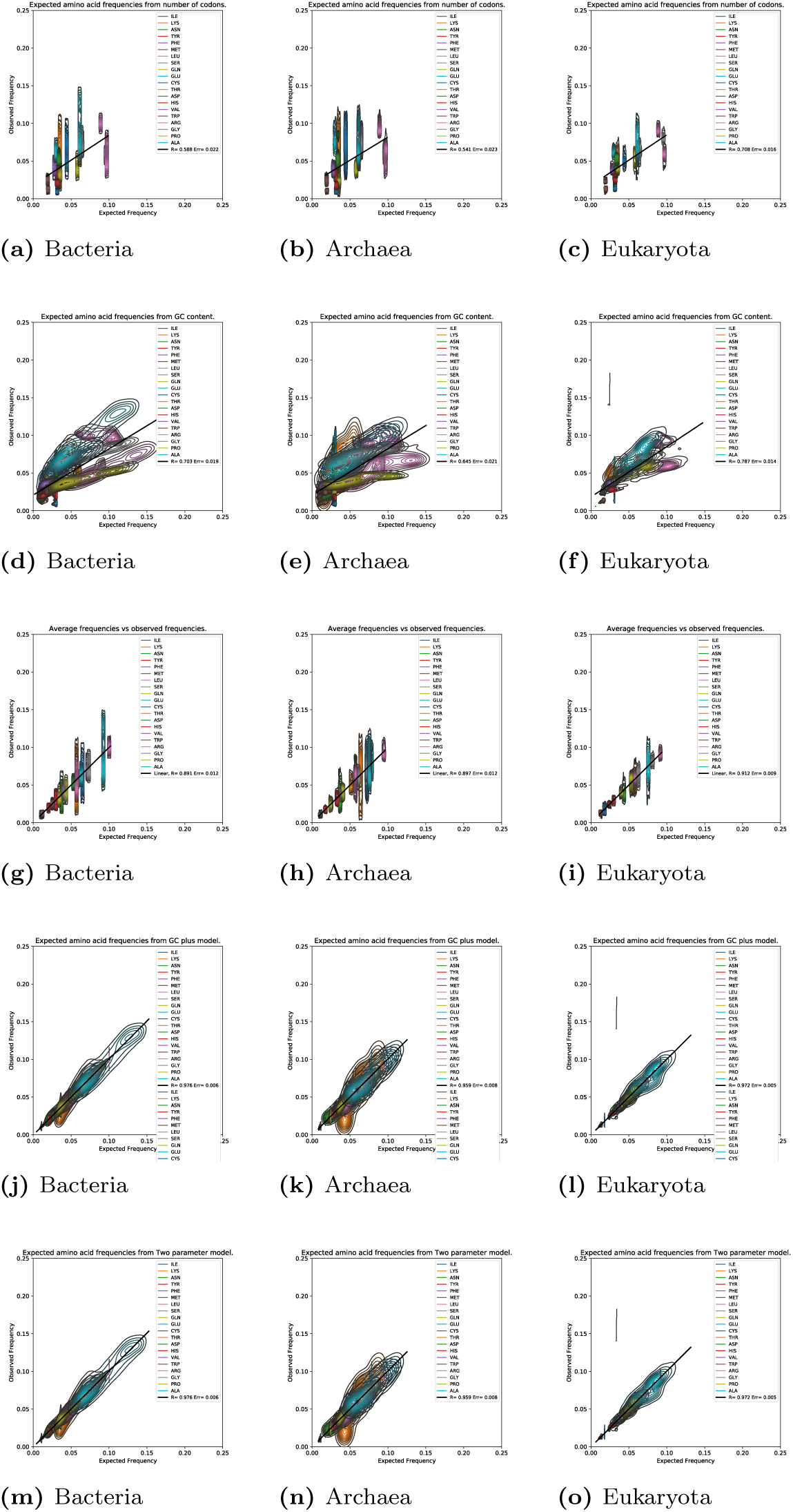
Correlation between expected and observed amino acid frequency for all amino acids over all genomes.

**Figure S2.**
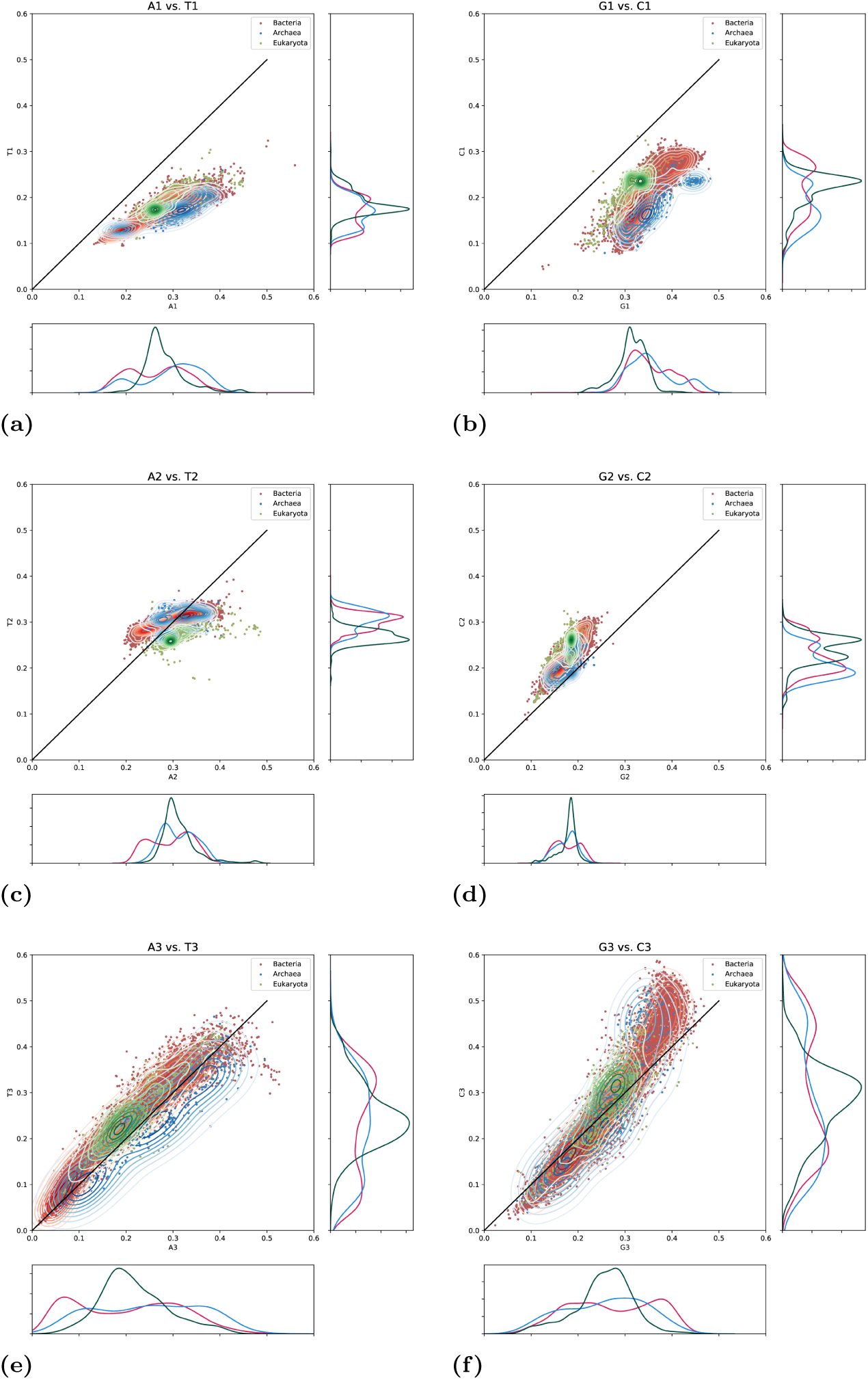
In position three perfect correlation, i.e. GC determines everything

**Figure S3.**
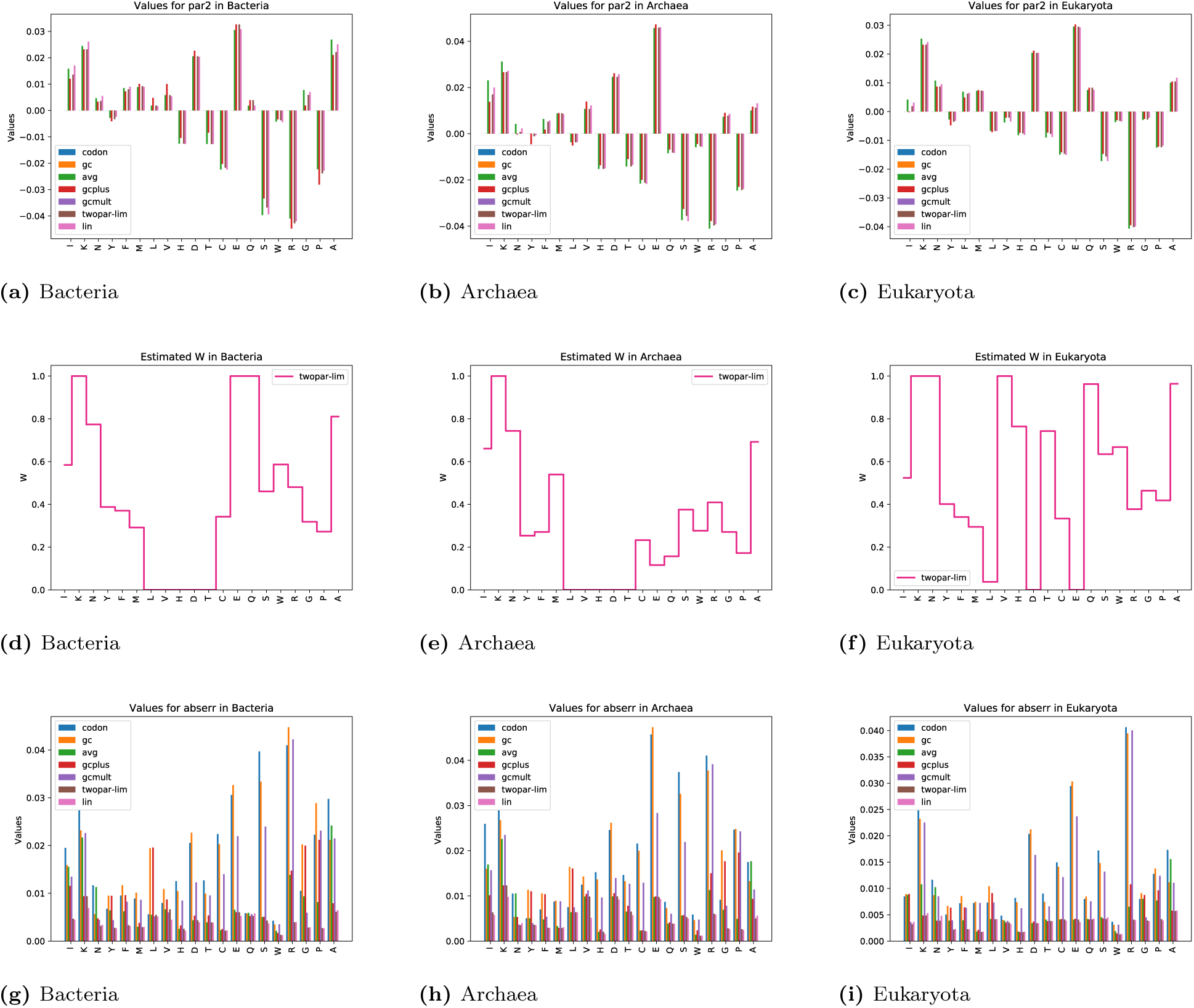
Combined plots from fitting parameters and absolute errors.

**Table S1.**
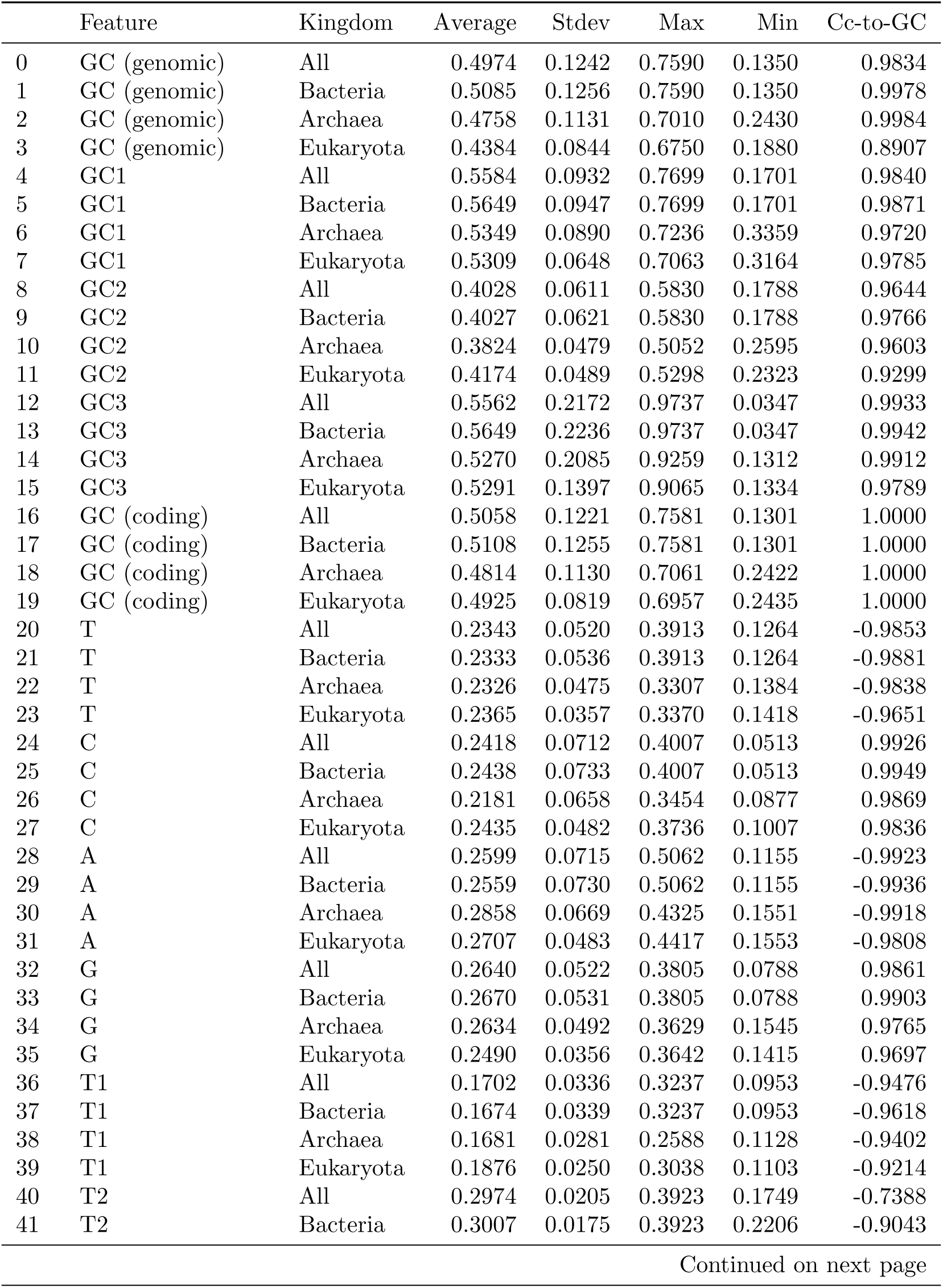
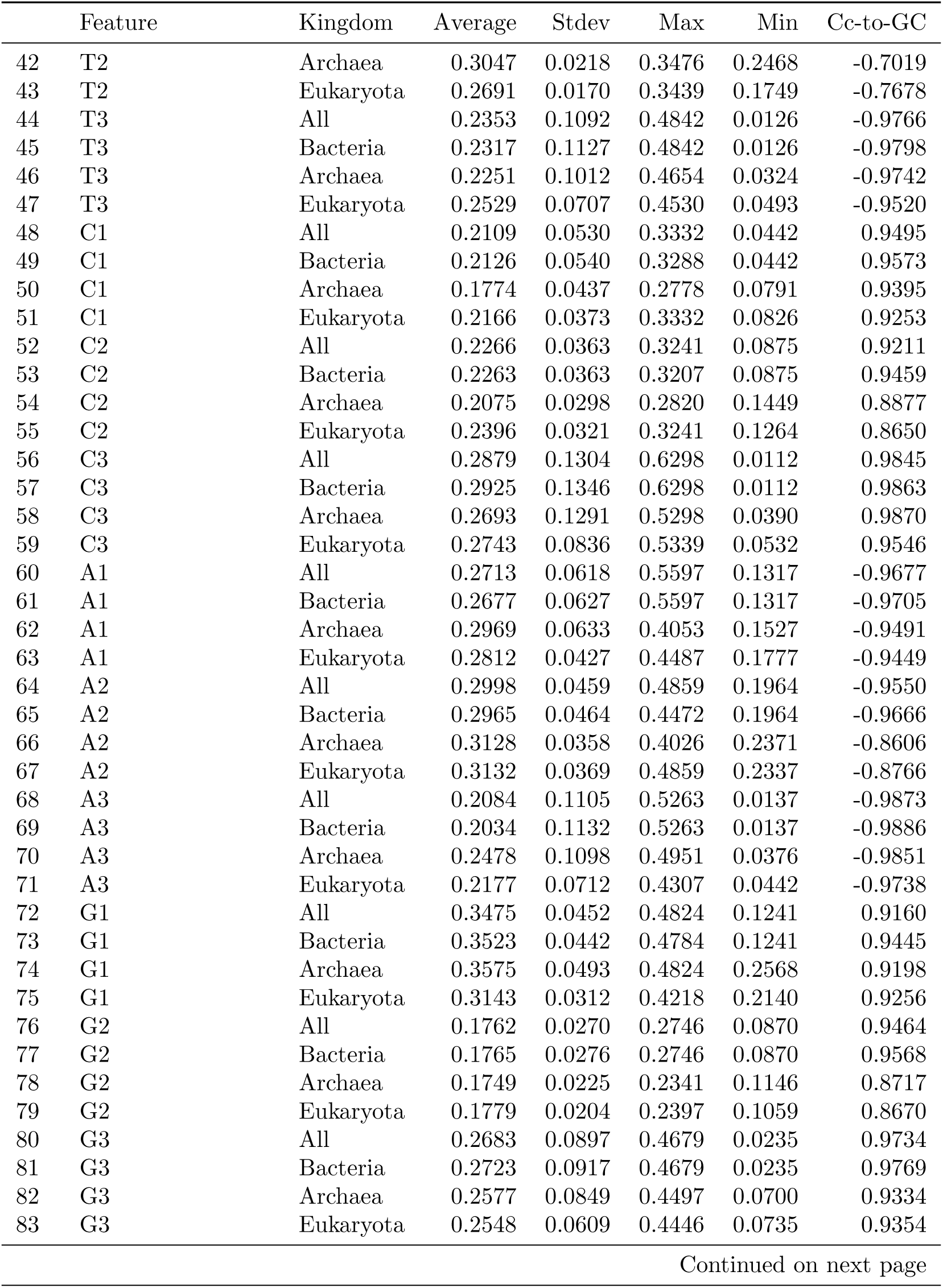
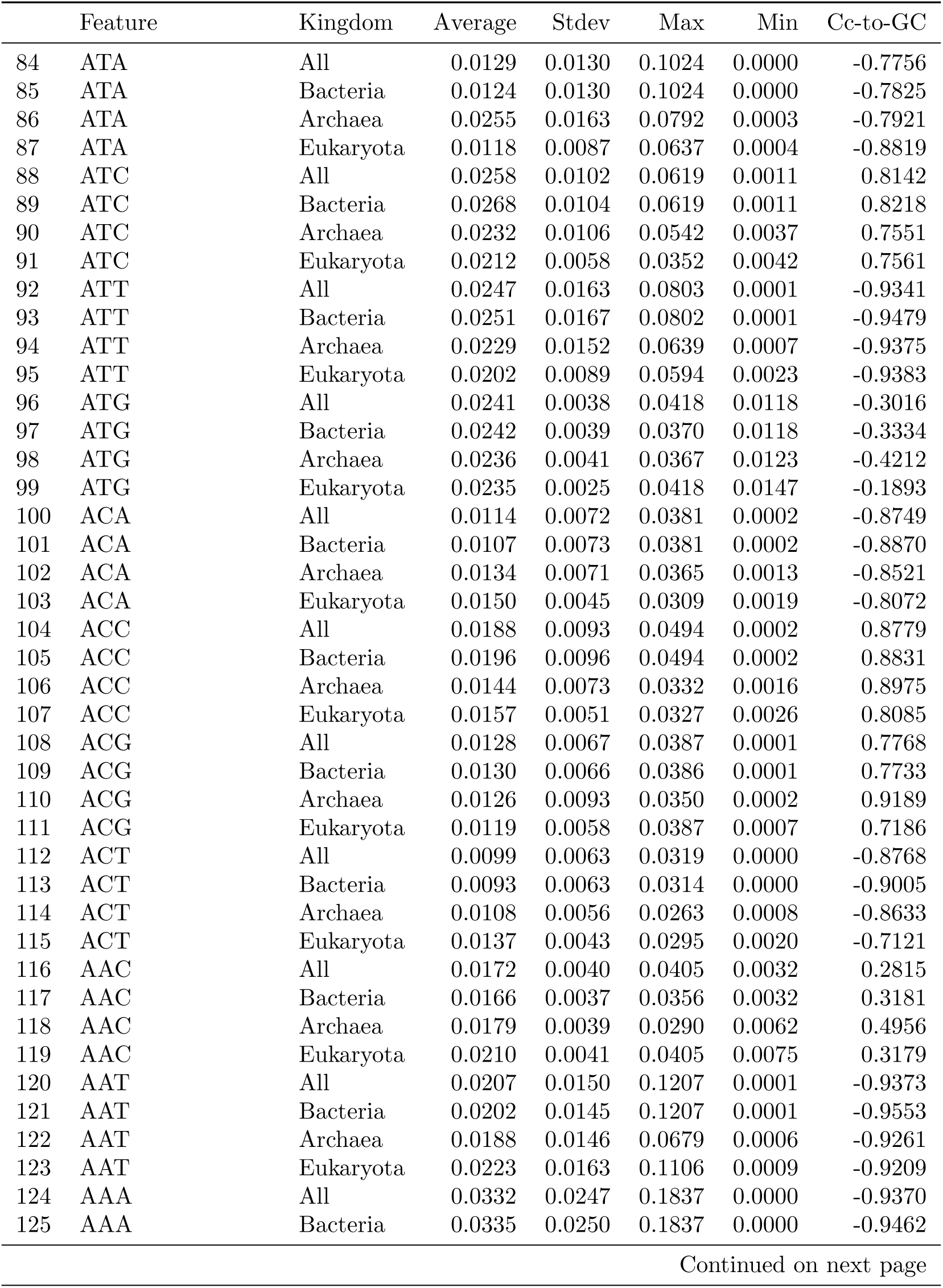
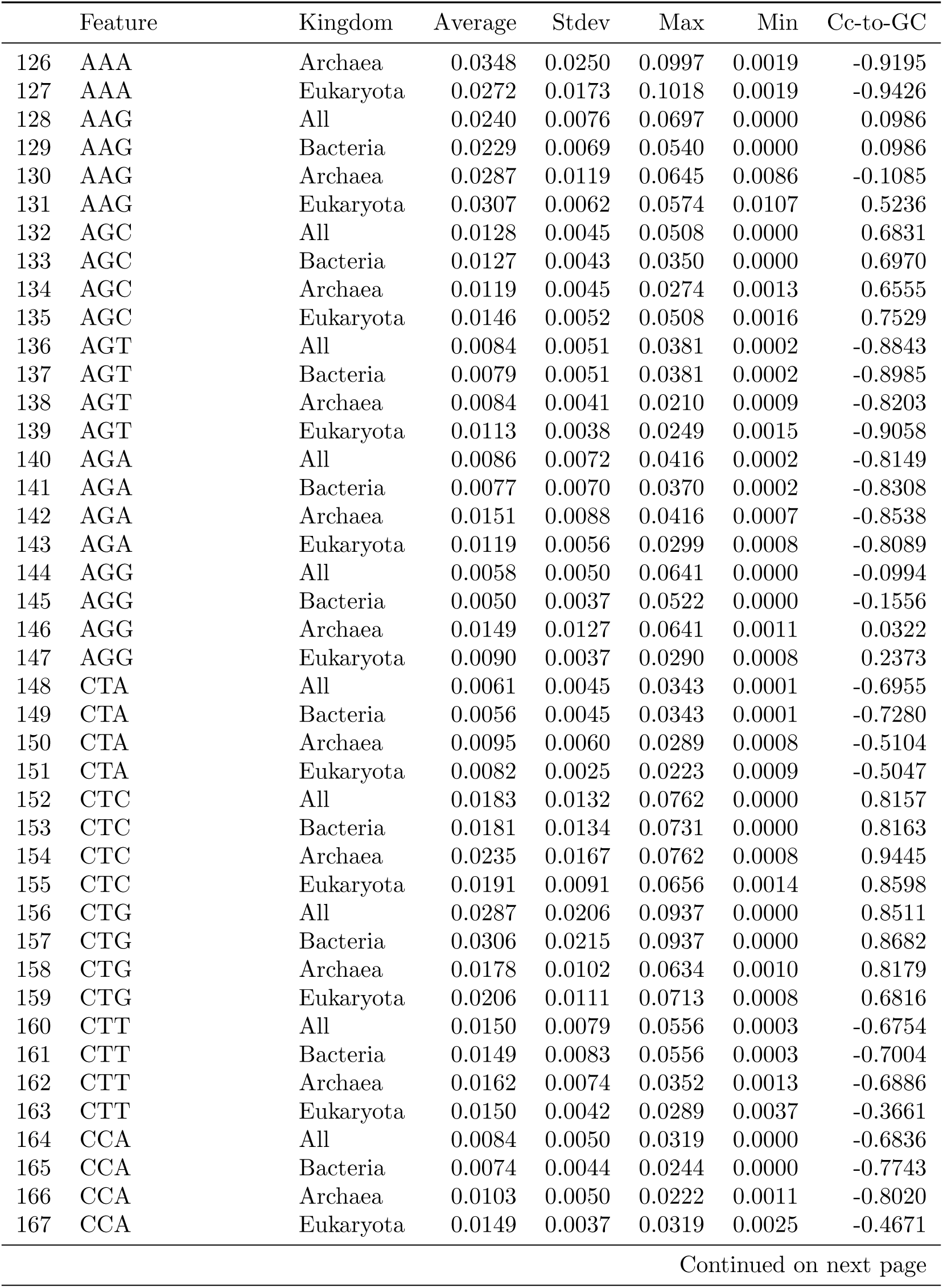
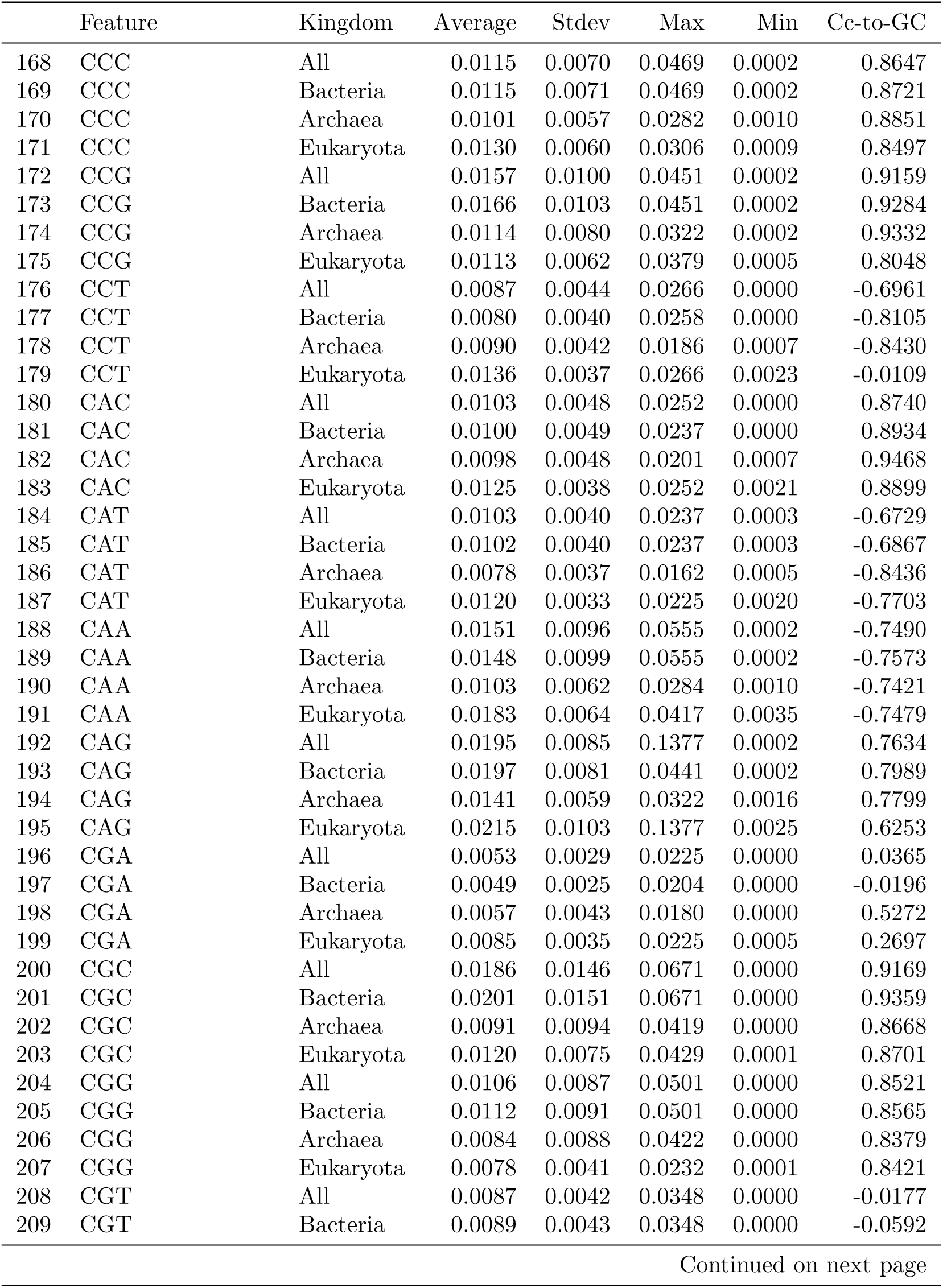
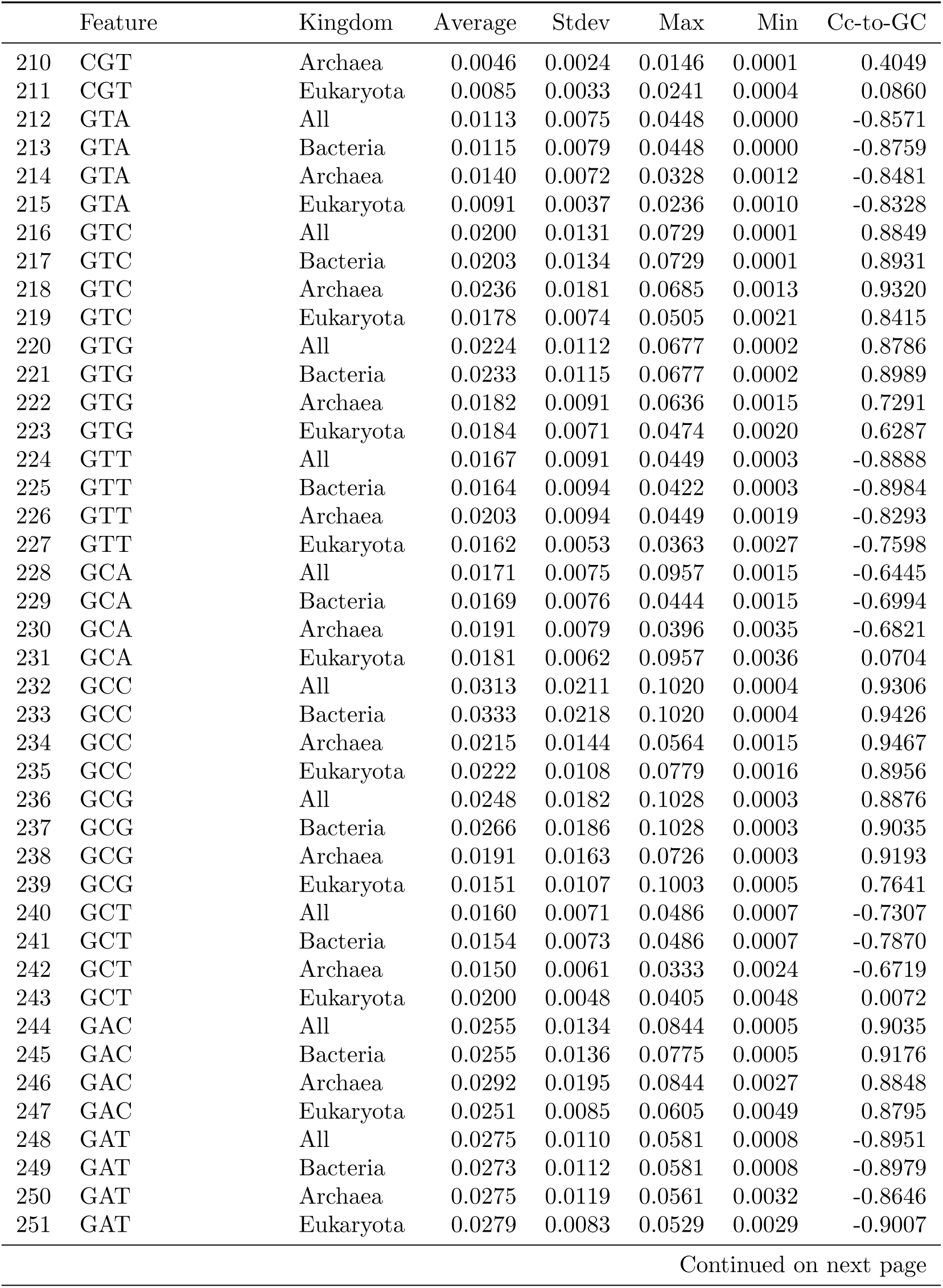
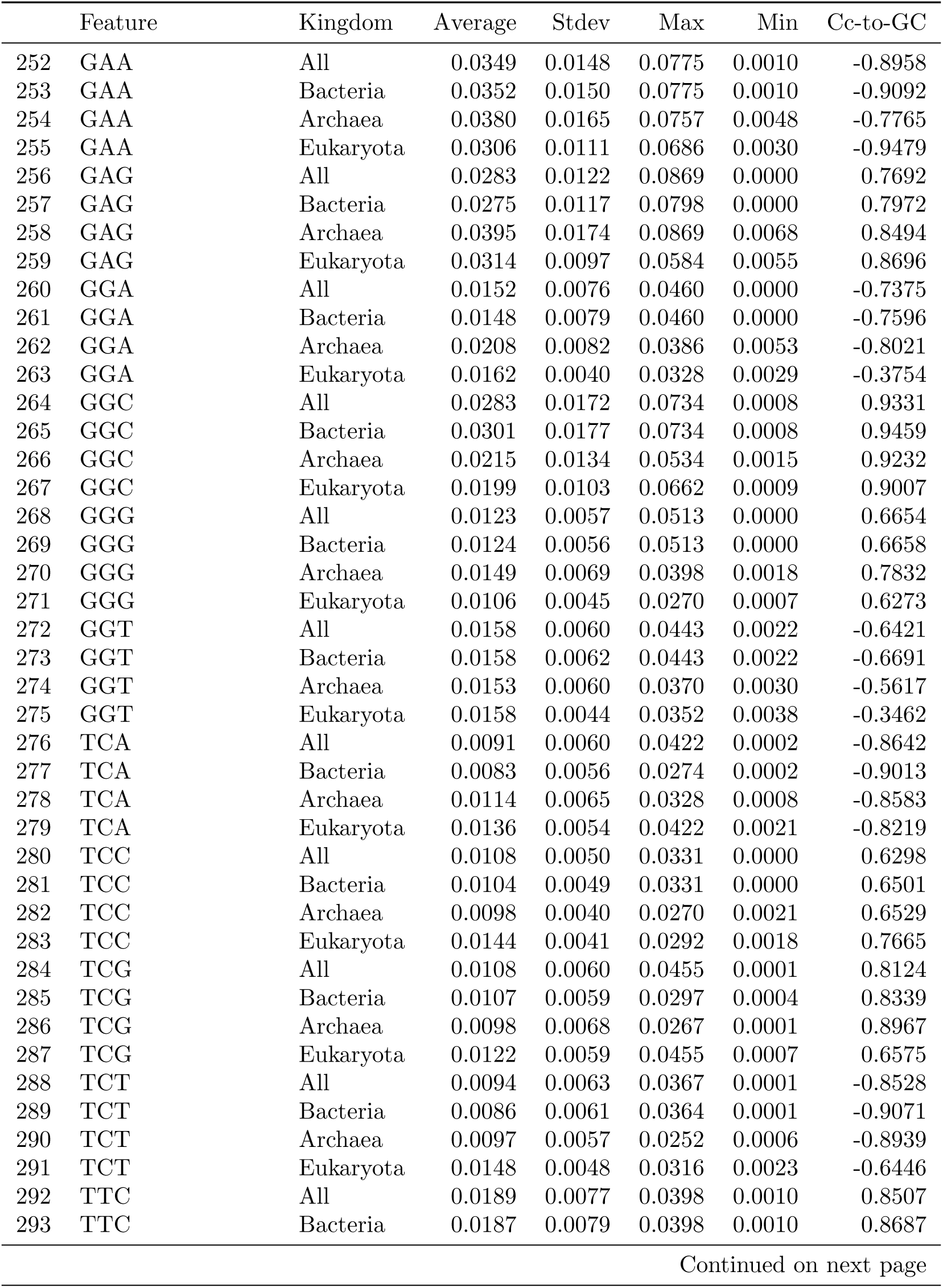
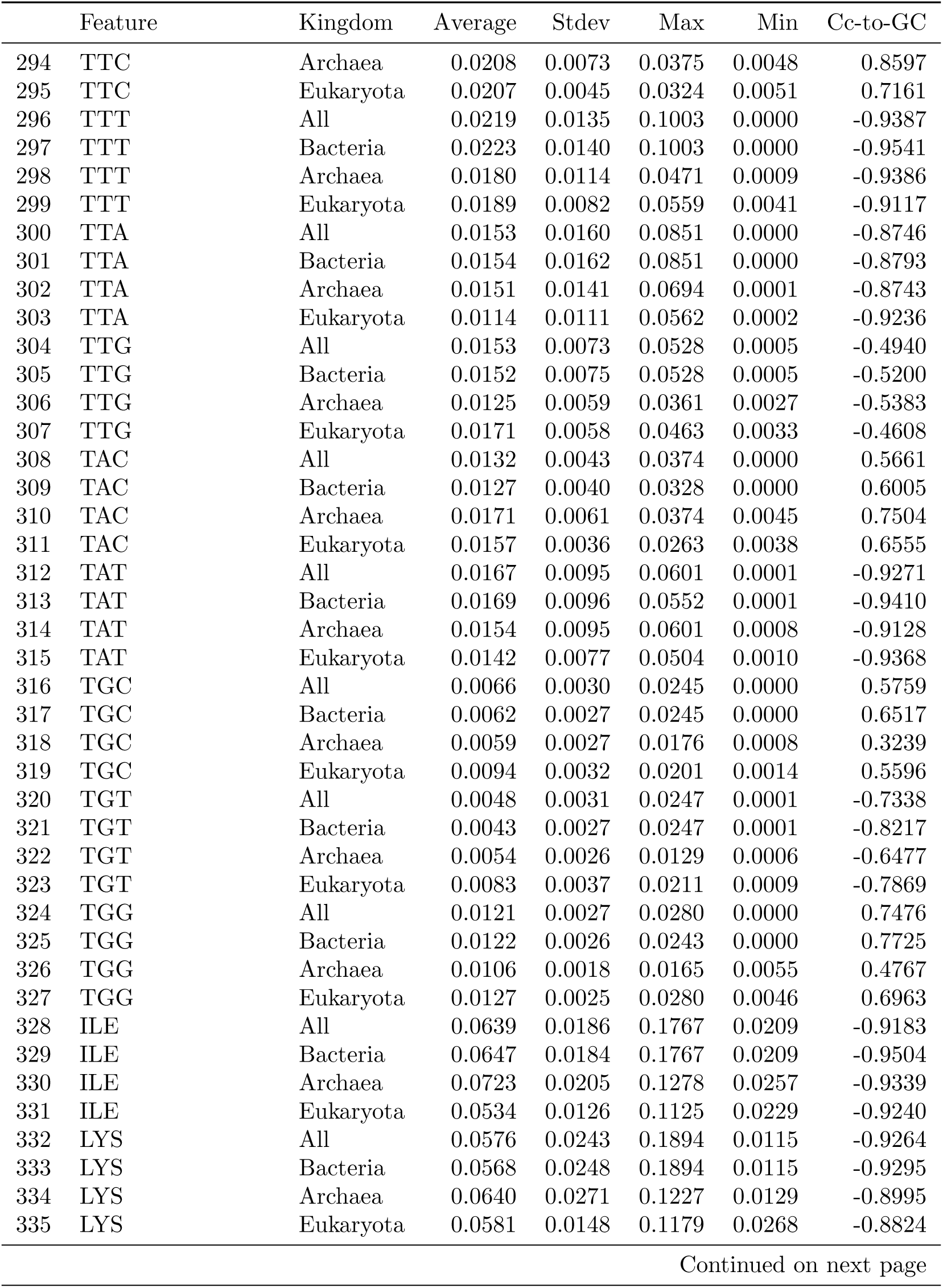
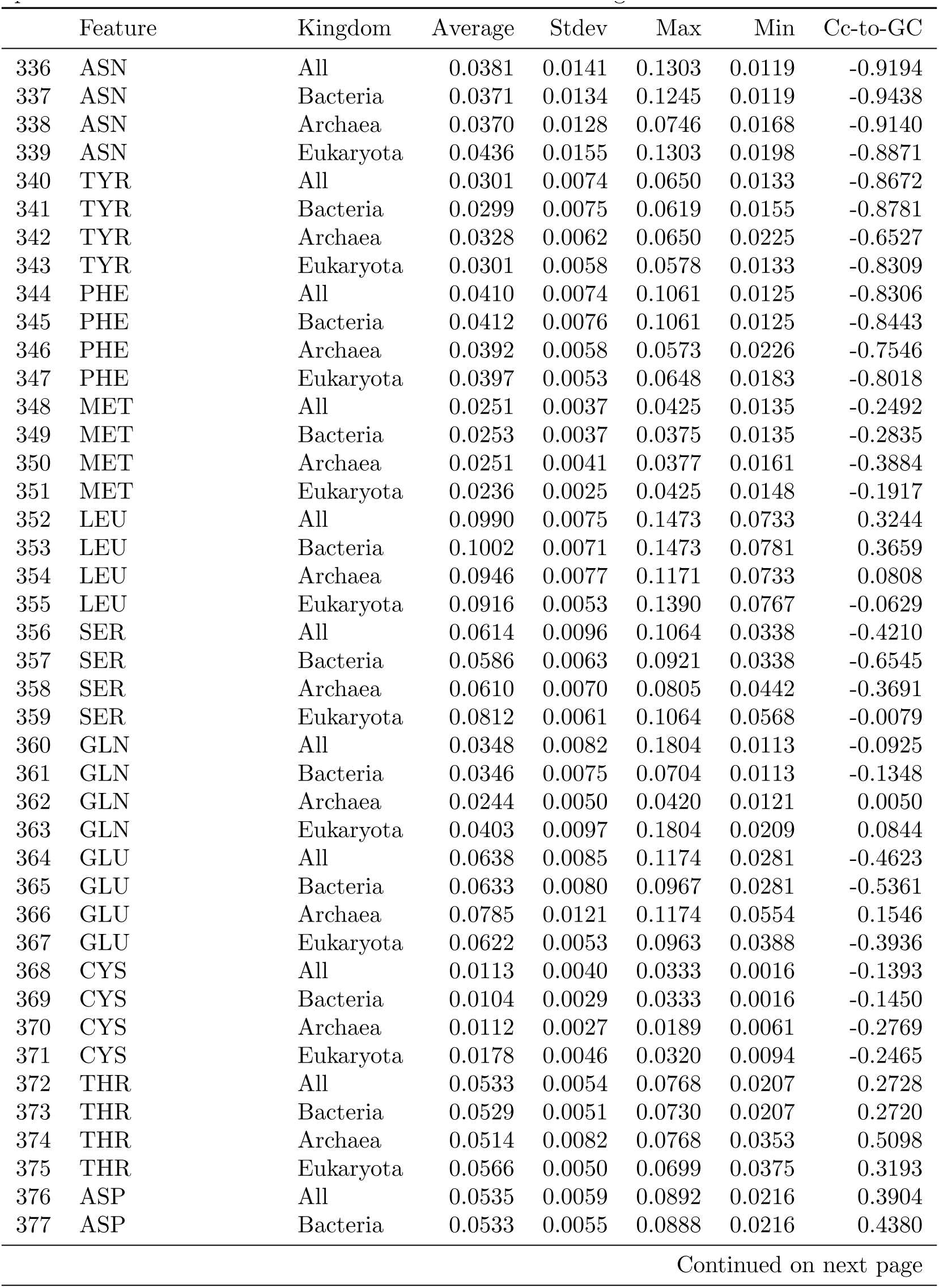
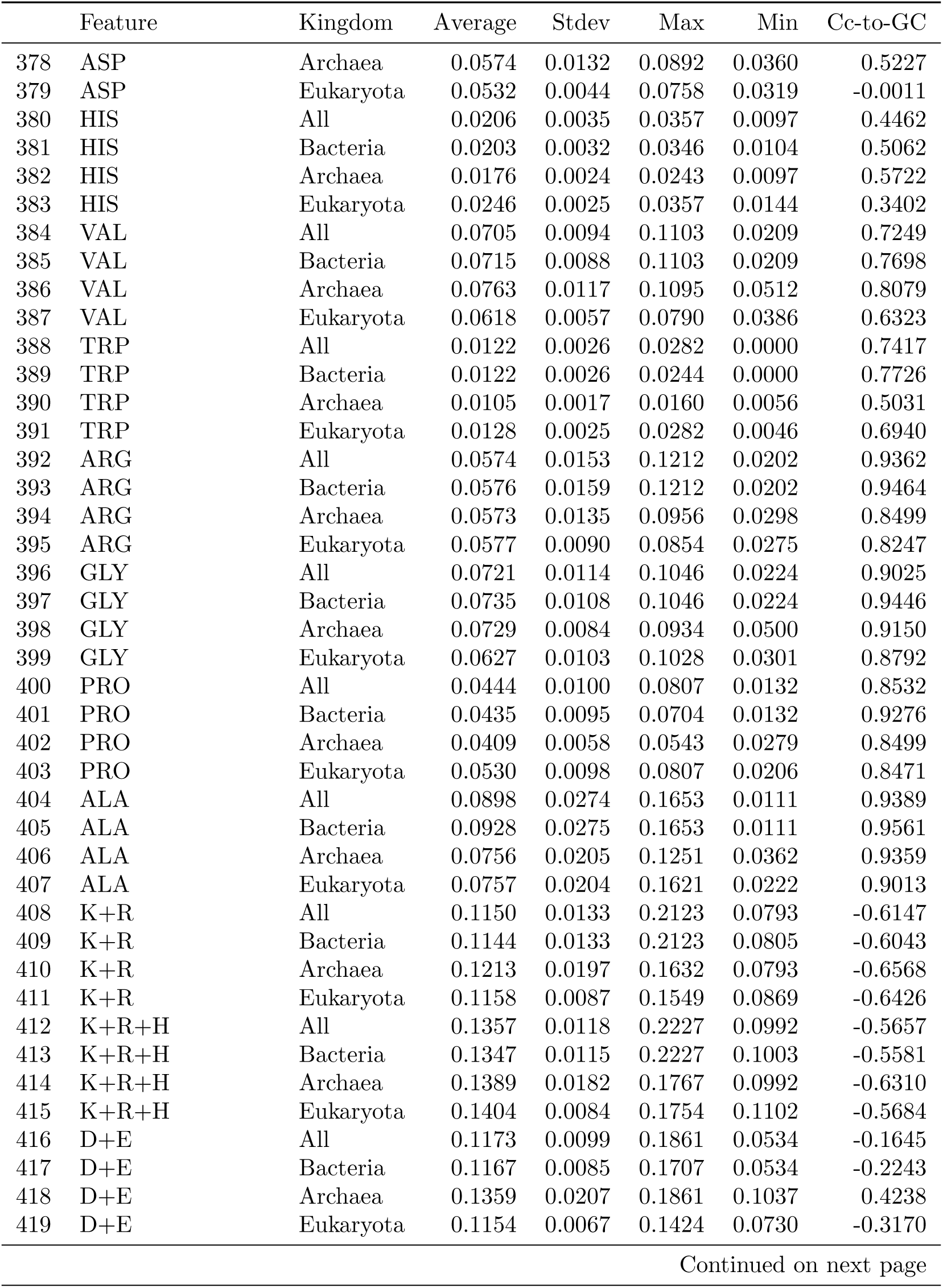
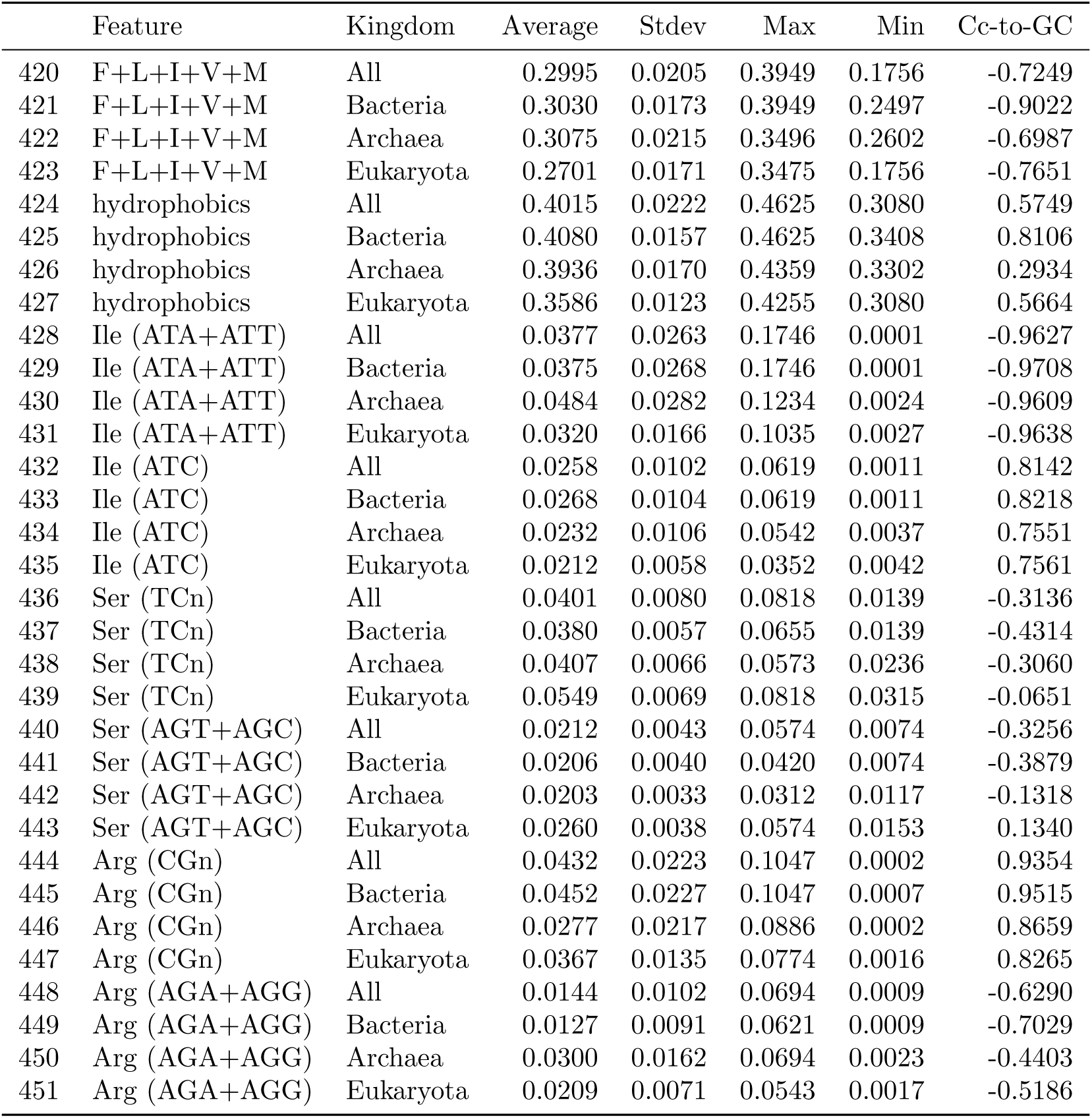
Summary table of all features in the three kingdoms. Average, standard deviation, max and min values are printed as well as the correlation coefficient with GCcoding.

|  | Feature | Kingdom | Average | Stdev | Max | Min | Cc-to-GC |
| --- | --- | --- | --- | --- | --- | --- | --- |
| 0 | GC (genomic) | All | 0.4974 | 0.1242 | 0.7590 | 0.1350 | 0.9834 |
| 1 | GC (genomic) | Bacteria | 0.5085 | 0.1256 | 0.7590 | 0.1350 | 0.9978 |
| 2 | GC (genomic) | Archaea | 0.4758 | 0.1131 | 0.7010 | 0.2430 | 0.9984 |
| 3 | GC (genomic) | Eukaryota | 0.4384 | 0.0844 | 0.6750 | 0.1880 | 0.8907 |
| 4 | GC1 | All | 0.5584 | 0.0932 | 0.7699 | 0.1701 | 0.9840 |
| 5 | GC1 | Bacteria | 0.5649 | 0.0947 | 0.7699 | 0.1701 | 0.9871 |
| 6 | GC1 | Archaea | 0.5349 | 0.0890 | 0.7236 | 0.3359 | 0.9720 |
| 7 | GC1 | Eukaryota | 0.5309 | 0.0648 | 0.7063 | 0.3164 | 0.9785 |
| 8 | GC2 | All | 0.4028 | 0.0611 | 0.5830 | 0.1788 | 0.9644 |
| 9 | GC2 | Bacteria | 0.4027 | 0.0621 | 0.5830 | 0.1788 | 0.9766 |
| 10 | GC2 | Archaea | 0.3824 | 0.0479 | 0.5052 | 0.2595 | 0.9603 |
| 11 | GC2 | Eukaryota | 0.4174 | 0.0489 | 0.5298 | 0.2323 | 0.9299 |
| 12 | GC3 | All | 0.5562 | 0.2172 | 0.9737 | 0.0347 | 0.9933 |
| 13 | GC3 | Bacteria | 0.5649 | 0.2236 | 0.9737 | 0.0347 | 0.9942 |
| 14 | GC3 | Archaea | 0.5270 | 0.2085 | 0.9259 | 0.1312 | 0.9912 |
| 15 | GC3 | Eukaryota | 0.5291 | 0.1397 | 0.9065 | 0.1334 | 0.9789 |
| 16 | GC (coding) | All | 0.5058 | 0.1221 | 0.7581 | 0.1301 | 1.0000 |
| 17 | GC (coding) | Bacteria | 0.5108 | 0.1255 | 0.7581 | 0.1301 | 1.0000 |
| 18 | GC (coding) | Archaea | 0.4814 | 0.1130 | 0.7061 | 0.2422 | 1.0000 |
| 19 | GC (coding) | Eukaryota | 0.4925 | 0.0819 | 0.6957 | 0.2435 | 1.0000 |
| 20 | T | All | 0.2343 | 0.0520 | 0.3913 | 0.1264 | -0.9853 |
| 21 | T | Bacteria | 0.2333 | 0.0536 | 0.3913 | 0.1264 | -0.9881 |
| 22 | T | Archaea | 0.2326 | 0.0475 | 0.3307 | 0.1384 | -0.9838 |
| 23 | T | Eukaryota | 0.2365 | 0.0357 | 0.3370 | 0.1418 | -0.9651 |
| 24 | C | All | 0.2418 | 0.0712 | 0.4007 | 0.0513 | 0.9926 |
| 25 | C | Bacteria | 0.2438 | 0.0733 | 0.4007 | 0.0513 | 0.9949 |
| 26 | C | Archaea | 0.2181 | 0.0658 | 0.3454 | 0.0877 | 0.9869 |
| 27 | C | Eukaryota | 0.2435 | 0.0482 | 0.3736 | 0.1007 | 0.9836 |
| 28 | A | All | 0.2599 | 0.0715 | 0.5062 | 0.1155 | -0.9923 |
| 29 | A | Bacteria | 0.2559 | 0.0730 | 0.5062 | 0.1155 | -0.9936 |
| 30 | A | Archaea | 0.2858 | 0.0669 | 0.4325 | 0.1551 | -0.9918 |
| 31 | A | Eukaryota | 0.2707 | 0.0483 | 0.4417 | 0.1553 | -0.9808 |
| 32 | G | All | 0.2640 | 0.0522 | 0.3805 | 0.0788 | 0.9861 |
| 33 | G | Bacteria | 0.2670 | 0.0531 | 0.3805 | 0.0788 | 0.9903 |
| 34 | G | Archaea | 0.2634 | 0.0492 | 0.3629 | 0.1545 | 0.9765 |
| 35 | G | Eukaryota | 0.2490 | 0.0356 | 0.3642 | 0.1415 | 0.9697 |
| 36 | T1 | All | 0.1702 | 0.0336 | 0.3237 | 0.0953 | -0.9476 |
| 37 | T1 | Bacteria | 0.1674 | 0.0339 | 0.3237 | 0.0953 | -0.9618 |
| 38 | T1 | Archaea | 0.1681 | 0.0281 | 0.2588 | 0.1128 | -0.9402 |
| 39 | T1 | Eukaryota | 0.1876 | 0.0250 | 0.3038 | 0.1103 | -0.9214 |
| 40 | T2 | All | 0.2974 | 0.0205 | 0.3923 | 0.1749 | -0.7388 |
| 41 | T2 | Bacteria | 0.3007 | 0.0175 | 0.3923 | 0.2206 | -0.9043 |
| 42 | T2 | Archaea | 0.3047 | 0.0218 | 0.3476 | 0.2468 | -0.7019 |
| 43 | T2 | Eukaryota | 0.2691 | 0.0170 | 0.3439 | 0.1749 | -0.7678 |
| 44 | T3 | All | 0.2353 | 0.1092 | 0.4842 | 0.0126 | -0.9766 |
| 45 | T3 | Bacteria | 0.2317 | 0.1127 | 0.4842 | 0.0126 | -0.9798 |
| 46 | T3 | Archaea | 0.2251 | 0.1012 | 0.4654 | 0.0324 | -0.9742 |
| 47 | T3 | Eukaryota | 0.2529 | 0.0707 | 0.4530 | 0.0493 | -0.9520 |
| 48 | C1 | All | 0.2109 | 0.0530 | 0.3332 | 0.0442 | 0.9495 |
| 49 | C1 | Bacteria | 0.2126 | 0.0540 | 0.3288 | 0.0442 | 0.9573 |
| 50 | C1 | Archaea | 0.1774 | 0.0437 | 0.2778 | 0.0791 | 0.9395 |
| 51 | C1 | Eukaryota | 0.2166 | 0.0373 | 0.3332 | 0.0826 | 0.9253 |
| 52 | C2 | All | 0.2266 | 0.0363 | 0.3241 | 0.0875 | 0.9211 |
| 53 | C2 | Bacteria | 0.2263 | 0.0363 | 0.3207 | 0.0875 | 0.9459 |
| 54 | C2 | Archaea | 0.2075 | 0.0298 | 0.2820 | 0.1449 | 0.8877 |
| 55 | C2 | Eukaryota | 0.2396 | 0.0321 | 0.3241 | 0.1264 | 0.8650 |
| 56 | C3 | All | 0.2879 | 0.1304 | 0.6298 | 0.0112 | 0.9845 |
| 57 | C3 | Bacteria | 0.2925 | 0.1346 | 0.6298 | 0.0112 | 0.9863 |
| 58 | C3 | Archaea | 0.2693 | 0.1291 | 0.5298 | 0.0390 | 0.9870 |
| 59 | C3 | Eukaryota | 0.2743 | 0.0836 | 0.5339 | 0.0532 | 0.9546 |
| 60 | A1 | All | 0.2713 | 0.0618 | 0.5597 | 0.1317 | -0.9677 |
| 61 | A1 | Bacteria | 0.2677 | 0.0627 | 0.5597 | 0.1317 | -0.9705 |
| 62 | A1 | Archaea | 0.2969 | 0.0633 | 0.4053 | 0.1527 | -0.9491 |
| 63 | A1 | Eukaryota | 0.2812 | 0.0427 | 0.4487 | 0.1777 | -0.9449 |
| 64 | A2 | All | 0.2998 | 0.0459 | 0.4859 | 0.1964 | -0.9550 |
| 65 | A2 | Bacteria | 0.2965 | 0.0464 | 0.4472 | 0.1964 | -0.9666 |
| 66 | A2 | Archaea | 0.3128 | 0.0358 | 0.4026 | 0.2371 | -0.8606 |
| 67 | A2 | Eukaryota | 0.3132 | 0.0369 | 0.4859 | 0.2337 | -0.8766 |
| 68 | A3 | All | 0.2084 | 0.1105 | 0.5263 | 0.0137 | -0.9873 |
| 69 | A3 | Bacteria | 0.2034 | 0.1132 | 0.5263 | 0.0137 | -0.9886 |
| 70 | A3 | Archaea | 0.2478 | 0.1098 | 0.4951 | 0.0376 | -0.9851 |
| 71 | A3 | Eukaryota | 0.2177 | 0.0712 | 0.4307 | 0.0442 | -0.9738 |
| 72 | G1 | All | 0.3475 | 0.0452 | 0.4824 | 0.1241 | 0.9160 |
| 73 | G1 | Bacteria | 0.3523 | 0.0442 | 0.4784 | 0.1241 | 0.9445 |
| 74 | G1 | Archaea | 0.3575 | 0.0493 | 0.4824 | 0.2568 | 0.9198 |
| 75 | G1 | Eukaryota | 0.3143 | 0.0312 | 0.4218 | 0.2140 | 0.9256 |
| 76 | G2 | All | 0.1762 | 0.0270 | 0.2746 | 0.0870 | 0.9464 |
| 77 | G2 | Bacteria | 0.1765 | 0.0276 | 0.2746 | 0.0870 | 0.9568 |
| 78 | G2 | Archaea | 0.1749 | 0.0225 | 0.2341 | 0.1146 | 0.8717 |
| 79 | G2 | Eukaryota | 0.1779 | 0.0204 | 0.2397 | 0.1059 | 0.8670 |
| 80 | G3 | All | 0.2683 | 0.0897 | 0.4679 | 0.0235 | 0.9734 |
| 81 | G3 | Bacteria | 0.2723 | 0.0917 | 0.4679 | 0.0235 | 0.9769 |
| 82 | G3 | Archaea | 0.2577 | 0.0849 | 0.4497 | 0.0700 | 0.9334 |
| 83 | G3 | Eukaryota | 0.2548 | 0.0609 | 0.4446 | 0.0735 | 0.9354 |
| 84 | ATA | All | 0.0129 | 0.0130 | 0.1024 | 0.0000 | -0.7756 |
| 85 | ATA | Bacteria | 0.0124 | 0.0130 | 0.1024 | 0.0000 | -0.7825 |
| 86 | ATA | Archaea | 0.0255 | 0.0163 | 0.0792 | 0.0003 | -0.7921 |
| 87 | ATA | Eukaryota | 0.0118 | 0.0087 | 0.0637 | 0.0004 | -0.8819 |
| 88 | ATC | All | 0.0258 | 0.0102 | 0.0619 | 0.0011 | 0.8142 |
| 89 | ATC | Bacteria | 0.0268 | 0.0104 | 0.0619 | 0.0011 | 0.8218 |
| 90 | ATC | Archaea | 0.0232 | 0.0106 | 0.0542 | 0.0037 | 0.7551 |
| 91 | ATC | Eukaryota | 0.0212 | 0.0058 | 0.0352 | 0.0042 | 0.7561 |
| 92 | ATT | All | 0.0247 | 0.0163 | 0.0803 | 0.0001 | -0.9341 |
| 93 | ATT | Bacteria | 0.0251 | 0.0167 | 0.0802 | 0.0001 | -0.9479 |
| 94 | ATT | Archaea | 0.0229 | 0.0152 | 0.0639 | 0.0007 | -0.9375 |
| 95 | ATT | Eukaryota | 0.0202 | 0.0089 | 0.0594 | 0.0023 | -0.9383 |
| 96 | ATG | All | 0.0241 | 0.0038 | 0.0418 | 0.0118 | -0.3016 |
| 97 | ATG | Bacteria | 0.0242 | 0.0039 | 0.0370 | 0.0118 | -0.3334 |
| 98 | ATG | Archaea | 0.0236 | 0.0041 | 0.0367 | 0.0123 | -0.4212 |
| 99 | ATG | Eukaryota | 0.0235 | 0.0025 | 0.0418 | 0.0147 | -0.1893 |
| 100 | ACA | All | 0.0114 | 0.0072 | 0.0381 | 0.0002 | -0.8749 |
| 101 | ACA | Bacteria | 0.0107 | 0.0073 | 0.0381 | 0.0002 | -0.8870 |
| 102 | ACA | Archaea | 0.0134 | 0.0071 | 0.0365 | 0.0013 | -0.8521 |
| 103 | ACA | Eukaryota | 0.0150 | 0.0045 | 0.0309 | 0.0019 | -0.8072 |
| 104 | ACC | All | 0.0188 | 0.0093 | 0.0494 | 0.0002 | 0.8779 |
| 105 | ACC | Bacteria | 0.0196 | 0.0096 | 0.0494 | 0.0002 | 0.8831 |
| 106 | ACC | Archaea | 0.0144 | 0.0073 | 0.0332 | 0.0016 | 0.8975 |
| 107 | ACC | Eukaryota | 0.0157 | 0.0051 | 0.0327 | 0.0026 | 0.8085 |
| 108 | ACG | All | 0.0128 | 0.0067 | 0.0387 | 0.0001 | 0.7768 |
| 109 | ACG | Bacteria | 0.0130 | 0.0066 | 0.0386 | 0.0001 | 0.7733 |
| 110 | ACG | Archaea | 0.0126 | 0.0093 | 0.0350 | 0.0002 | 0.9189 |
| 111 | ACG | Eukaryota | 0.0119 | 0.0058 | 0.0387 | 0.0007 | 0.7186 |
| 112 | ACT | All | 0.0099 | 0.0063 | 0.0319 | 0.0000 | -0.8768 |
| 113 | ACT | Bacteria | 0.0093 | 0.0063 | 0.0314 | 0.0000 | -0.9005 |
| 114 | ACT | Archaea | 0.0108 | 0.0056 | 0.0263 | 0.0008 | -0.8633 |
| 115 | ACT | Eukaryota | 0.0137 | 0.0043 | 0.0295 | 0.0020 | -0.7121 |
| 116 | AAC | All | 0.0172 | 0.0040 | 0.0405 | 0.0032 | 0.2815 |
| 117 | AAC | Bacteria | 0.0166 | 0.0037 | 0.0356 | 0.0032 | 0.3181 |
| 118 | AAC | Archaea | 0.0179 | 0.0039 | 0.0290 | 0.0062 | 0.4956 |
| 119 | AAC | Eukaryota | 0.0210 | 0.0041 | 0.0405 | 0.0075 | 0.3179 |
| 120 | AAT | All | 0.0207 | 0.0150 | 0.1207 | 0.0001 | -0.9373 |
| 121 | AAT | Bacteria | 0.0202 | 0.0145 | 0.1207 | 0.0001 | -0.9553 |
| 122 | AAT | Archaea | 0.0188 | 0.0146 | 0.0679 | 0.0006 | -0.9261 |
| 123 | AAT | Eukaryota | 0.0223 | 0.0163 | 0.1106 | 0.0009 | -0.9209 |
| 124 | AAA | All | 0.0332 | 0.0247 | 0.1837 | 0.0000 | -0.9370 |
| 125 | AAA | Bacteria | 0.0335 | 0.0250 | 0.1837 | 0.0000 | -0.9462 |
| 126 | AAA | Archaea | 0.0348 | 0.0250 | 0.0997 | 0.0019 | -0.9195 |
| 127 | AAA | Eukaryota | 0.0272 | 0.0173 | 0.1018 | 0.0019 | -0.9426 |
| 128 | AAG | All | 0.0240 | 0.0076 | 0.0697 | 0.0000 | 0.0986 |
| 129 | AAG | Bacteria | 0.0229 | 0.0069 | 0.0540 | 0.0000 | 0.0986 |
| 130 | AAG | Archaea | 0.0287 | 0.0119 | 0.0645 | 0.0086 | -0.1085 |
| 131 | AAG | Eukaryota | 0.0307 | 0.0062 | 0.0574 | 0.0107 | 0.5236 |
| 132 | AGC | All | 0.0128 | 0.0045 | 0.0508 | 0.0000 | 0.6831 |
| 133 | AGC | Bacteria | 0.0127 | 0.0043 | 0.0350 | 0.0000 | 0.6970 |
| 134 | AGC | Archaea | 0.0119 | 0.0045 | 0.0274 | 0.0013 | 0.6555 |
| 135 | AGC | Eukaryota | 0.0146 | 0.0052 | 0.0508 | 0.0016 | 0.7529 |
| 136 | AGT | All | 0.0084 | 0.0051 | 0.0381 | 0.0002 | -0.8843 |
| 137 | AGT | Bacteria | 0.0079 | 0.0051 | 0.0381 | 0.0002 | -0.8985 |
| 138 | AGT | Archaea | 0.0084 | 0.0041 | 0.0210 | 0.0009 | -0.8203 |
| 139 | AGT | Eukaryota | 0.0113 | 0.0038 | 0.0249 | 0.0015 | -0.9058 |
| 140 | AGA | All | 0.0086 | 0.0072 | 0.0416 | 0.0002 | -0.8149 |
| 141 | AGA | Bacteria | 0.0077 | 0.0070 | 0.0370 | 0.0002 | -0.8308 |
| 142 | AGA | Archaea | 0.0151 | 0.0088 | 0.0416 | 0.0007 | -0.8538 |
| 143 | AGA | Eukaryota | 0.0119 | 0.0056 | 0.0299 | 0.0008 | -0.8089 |
| 144 | AGG | All | 0.0058 | 0.0050 | 0.0641 | 0.0000 | -0.0994 |
| 145 | AGG | Bacteria | 0.0050 | 0.0037 | 0.0522 | 0.0000 | -0.1556 |
| 146 | AGG | Archaea | 0.0149 | 0.0127 | 0.0641 | 0.0011 | 0.0322 |
| 147 | AGG | Eukaryota | 0.0090 | 0.0037 | 0.0290 | 0.0008 | 0.2373 |
| 148 | CTA | All | 0.0061 | 0.0045 | 0.0343 | 0.0001 | -0.6955 |
| 149 | CTA | Bacteria | 0.0056 | 0.0045 | 0.0343 | 0.0001 | -0.7280 |
| 150 | CTA | Archaea | 0.0095 | 0.0060 | 0.0289 | 0.0008 | -0.5104 |
| 151 | CTA | Eukaryota | 0.0082 | 0.0025 | 0.0223 | 0.0009 | -0.5047 |
| 152 | CTC | All | 0.0183 | 0.0132 | 0.0762 | 0.0000 | 0.8157 |
| 153 | CTC | Bacteria | 0.0181 | 0.0134 | 0.0731 | 0.0000 | 0.8163 |
| 154 | CTC | Archaea | 0.0235 | 0.0167 | 0.0762 | 0.0008 | 0.9445 |
| 155 | CTC | Eukaryota | 0.0191 | 0.0091 | 0.0656 | 0.0014 | 0.8598 |
| 156 | CTG | All | 0.0287 | 0.0206 | 0.0937 | 0.0000 | 0.8511 |
| 157 | CTG | Bacteria | 0.0306 | 0.0215 | 0.0937 | 0.0000 | 0.8682 |
| 158 | CTG | Archaea | 0.0178 | 0.0102 | 0.0634 | 0.0010 | 0.8179 |
| 159 | CTG | Eukaryota | 0.0206 | 0.0111 | 0.0713 | 0.0008 | 0.6816 |
| 160 | CTT | All | 0.0150 | 0.0079 | 0.0556 | 0.0003 | -0.6754 |
| 161 | CTT | Bacteria | 0.0149 | 0.0083 | 0.0556 | 0.0003 | -0.7004 |
| 162 | CTT | Archaea | 0.0162 | 0.0074 | 0.0352 | 0.0013 | -0.6886 |
| 163 | CTT | Eukaryota | 0.0150 | 0.0042 | 0.0289 | 0.0037 | -0.3661 |
| 164 | CCA | All | 0.0084 | 0.0050 | 0.0319 | 0.0000 | -0.6836 |
| 165 | CCA | Bacteria | 0.0074 | 0.0044 | 0.0244 | 0.0000 | -0.7743 |
| 166 | CCA | Archaea | 0.0103 | 0.0050 | 0.0222 | 0.0011 | -0.8020 |
| 167 | CCA | Eukaryota | 0.0149 | 0.0037 | 0.0319 | 0.0025 | -0.4671 |
| 168 | CCC | All | 0.0115 | 0.0070 | 0.0469 | 0.0002 | 0.8647 |
| 169 | CCC | Bacteria | 0.0115 | 0.0071 | 0.0469 | 0.0002 | 0.8721 |
| 170 | CCC | Archaea | 0.0101 | 0.0057 | 0.0282 | 0.0010 | 0.8851 |
| 171 | CCC | Eukaryota | 0.0130 | 0.0060 | 0.0306 | 0.0009 | 0.8497 |
| 172 | CCG | All | 0.0157 | 0.0100 | 0.0451 | 0.0002 | 0.9159 |
| 173 | CCG | Bacteria | 0.0166 | 0.0103 | 0.0451 | 0.0002 | 0.9284 |
| 174 | CCG | Archaea | 0.0114 | 0.0080 | 0.0322 | 0.0002 | 0.9332 |
| 175 | CCG | Eukaryota | 0.0113 | 0.0062 | 0.0379 | 0.0005 | 0.8048 |
| 176 | CCT | All | 0.0087 | 0.0044 | 0.0266 | 0.0000 | -0.6961 |
| 177 | CCT | Bacteria | 0.0080 | 0.0040 | 0.0258 | 0.0000 | -0.8105 |
| 178 | CCT | Archaea | 0.0090 | 0.0042 | 0.0186 | 0.0007 | -0.8430 |
| 179 | CCT | Eukaryota | 0.0136 | 0.0037 | 0.0266 | 0.0023 | -0.0109 |
| 180 | CAC | All | 0.0103 | 0.0048 | 0.0252 | 0.0000 | 0.8740 |
| 181 | CAC | Bacteria | 0.0100 | 0.0049 | 0.0237 | 0.0000 | 0.8934 |
| 182 | CAC | Archaea | 0.0098 | 0.0048 | 0.0201 | 0.0007 | 0.9468 |
| 183 | CAC | Eukaryota | 0.0125 | 0.0038 | 0.0252 | 0.0021 | 0.8899 |
| 184 | CAT | All | 0.0103 | 0.0040 | 0.0237 | 0.0003 | -0.6729 |
| 185 | CAT | Bacteria | 0.0102 | 0.0040 | 0.0237 | 0.0003 | -0.6867 |
| 186 | CAT | Archaea | 0.0078 | 0.0037 | 0.0162 | 0.0005 | -0.8436 |
| 187 | CAT | Eukaryota | 0.0120 | 0.0033 | 0.0225 | 0.0020 | -0.7703 |
| 188 | CAA | All | 0.0151 | 0.0096 | 0.0555 | 0.0002 | -0.7490 |
| 189 | CAA | Bacteria | 0.0148 | 0.0099 | 0.0555 | 0.0002 | -0.7573 |
| 190 | CAA | Archaea | 0.0103 | 0.0062 | 0.0284 | 0.0010 | -0.7421 |
| 191 | CAA | Eukaryota | 0.0183 | 0.0064 | 0.0417 | 0.0035 | -0.7479 |
| 192 | CAG | All | 0.0195 | 0.0085 | 0.1377 | 0.0002 | 0.7634 |
| 193 | CAG | Bacteria | 0.0197 | 0.0081 | 0.0441 | 0.0002 | 0.7989 |
| 194 | CAG | Archaea | 0.0141 | 0.0059 | 0.0322 | 0.0016 | 0.7799 |
| 195 | CAG | Eukaryota | 0.0215 | 0.0103 | 0.1377 | 0.0025 | 0.6253 |
| 196 | CGA | All | 0.0053 | 0.0029 | 0.0225 | 0.0000 | 0.0365 |
| 197 | CGA | Bacteria | 0.0049 | 0.0025 | 0.0204 | 0.0000 | -0.0196 |
| 198 | CGA | Archaea | 0.0057 | 0.0043 | 0.0180 | 0.0000 | 0.5272 |
| 199 | CGA | Eukaryota | 0.0085 | 0.0035 | 0.0225 | 0.0005 | 0.2697 |
| 200 | CGC | All | 0.0186 | 0.0146 | 0.0671 | 0.0000 | 0.9169 |
| 201 | CGC | Bacteria | 0.0201 | 0.0151 | 0.0671 | 0.0000 | 0.9359 |
| 202 | CGC | Archaea | 0.0091 | 0.0094 | 0.0419 | 0.0000 | 0.8668 |
| 203 | CGC | Eukaryota | 0.0120 | 0.0075 | 0.0429 | 0.0001 | 0.8701 |
| 204 | CGG | All | 0.0106 | 0.0087 | 0.0501 | 0.0000 | 0.8521 |
| 205 | CGG | Bacteria | 0.0112 | 0.0091 | 0.0501 | 0.0000 | 0.8565 |
| 206 | CGG | Archaea | 0.0084 | 0.0088 | 0.0422 | 0.0000 | 0.8379 |
| 207 | CGG | Eukaryota | 0.0078 | 0.0041 | 0.0232 | 0.0001 | 0.8421 |
| 208 | CGT | All | 0.0087 | 0.0042 | 0.0348 | 0.0000 | -0.0177 |
| 209 | CGT | Bacteria | 0.0089 | 0.0043 | 0.0348 | 0.0000 | -0.0592 |
| 210 | CGT | Archaea | 0.0046 | 0.0024 | 0.0146 | 0.0001 | 0.4049 |
| 211 | CGT | Eukaryota | 0.0085 | 0.0033 | 0.0241 | 0.0004 | 0.0860 |
| 212 | GTA | All | 0.0113 | 0.0075 | 0.0448 | 0.0000 | -0.8571 |
| 213 | GTA | Bacteria | 0.0115 | 0.0079 | 0.0448 | 0.0000 | -0.8759 |
| 214 | GTA | Archaea | 0.0140 | 0.0072 | 0.0328 | 0.0012 | -0.8481 |
| 215 | GTA | Eukaryota | 0.0091 | 0.0037 | 0.0236 | 0.0010 | -0.8328 |
| 216 | GTC | All | 0.0200 | 0.0131 | 0.0729 | 0.0001 | 0.8849 |
| 217 | GTC | Bacteria | 0.0203 | 0.0134 | 0.0729 | 0.0001 | 0.8931 |
| 218 | GTC | Archaea | 0.0236 | 0.0181 | 0.0685 | 0.0013 | 0.9320 |
| 219 | GTC | Eukaryota | 0.0178 | 0.0074 | 0.0505 | 0.0021 | 0.8415 |
| 220 | GTG | All | 0.0224 | 0.0112 | 0.0677 | 0.0002 | 0.8786 |
| 221 | GTG | Bacteria | 0.0233 | 0.0115 | 0.0677 | 0.0002 | 0.8989 |
| 222 | GTG | Archaea | 0.0182 | 0.0091 | 0.0636 | 0.0015 | 0.7291 |
| 223 | GTG | Eukaryota | 0.0184 | 0.0071 | 0.0474 | 0.0020 | 0.6287 |
| 224 | GTT | All | 0.0167 | 0.0091 | 0.0449 | 0.0003 | -0.8888 |
| 225 | GTT | Bacteria | 0.0164 | 0.0094 | 0.0422 | 0.0003 | -0.8984 |
| 226 | GTT | Archaea | 0.0203 | 0.0094 | 0.0449 | 0.0019 | -0.8293 |
| 227 | GTT | Eukaryota | 0.0162 | 0.0053 | 0.0363 | 0.0027 | -0.7598 |
| 228 | GCA | All | 0.0171 | 0.0075 | 0.0957 | 0.0015 | -0.6445 |
| 229 | GCA | Bacteria | 0.0169 | 0.0076 | 0.0444 | 0.0015 | -0.6994 |
| 230 | GCA | Archaea | 0.0191 | 0.0079 | 0.0396 | 0.0035 | -0.6821 |
| 231 | GCA | Eukaryota | 0.0181 | 0.0062 | 0.0957 | 0.0036 | 0.0704 |
| 232 | GCC | All | 0.0313 | 0.0211 | 0.1020 | 0.0004 | 0.9306 |
| 233 | GCC | Bacteria | 0.0333 | 0.0218 | 0.1020 | 0.0004 | 0.9426 |
| 234 | GCC | Archaea | 0.0215 | 0.0144 | 0.0564 | 0.0015 | 0.9467 |
| 235 | GCC | Eukaryota | 0.0222 | 0.0108 | 0.0779 | 0.0016 | 0.8956 |
| 236 | GCG | All | 0.0248 | 0.0182 | 0.1028 | 0.0003 | 0.8876 |
| 237 | GCG | Bacteria | 0.0266 | 0.0186 | 0.1028 | 0.0003 | 0.9035 |
| 238 | GCG | Archaea | 0.0191 | 0.0163 | 0.0726 | 0.0003 | 0.9193 |
| 239 | GCG | Eukaryota | 0.0151 | 0.0107 | 0.1003 | 0.0005 | 0.7641 |
| 240 | GCT | All | 0.0160 | 0.0071 | 0.0486 | 0.0007 | -0.7307 |
| 241 | GCT | Bacteria | 0.0154 | 0.0073 | 0.0486 | 0.0007 | -0.7870 |
| 242 | GCT | Archaea | 0.0150 | 0.0061 | 0.0333 | 0.0024 | -0.6719 |
| 243 | GCT | Eukaryota | 0.0200 | 0.0048 | 0.0405 | 0.0048 | 0.0072 |
| 244 | GAC | All | 0.0255 | 0.0134 | 0.0844 | 0.0005 | 0.9035 |
| 245 | GAC | Bacteria | 0.0255 | 0.0136 | 0.0775 | 0.0005 | 0.9176 |
| 246 | GAC | Archaea | 0.0292 | 0.0195 | 0.0844 | 0.0027 | 0.8848 |
| 247 | GAC | Eukaryota | 0.0251 | 0.0085 | 0.0605 | 0.0049 | 0.8795 |
| 248 | GAT | All | 0.0275 | 0.0110 | 0.0581 | 0.0008 | -0.8951 |
| 249 | GAT | Bacteria | 0.0273 | 0.0112 | 0.0581 | 0.0008 | -0.8979 |
| 250 | GAT | Archaea | 0.0275 | 0.0119 | 0.0561 | 0.0032 | -0.8646 |
| 251 | GAT | Eukaryota | 0.0279 | 0.0083 | 0.0529 | 0.0029 | -0.9007 |
| 252 | GAA | All | 0.0349 | 0.0148 | 0.0775 | 0.0010 | -0.8958 |
| 253 | GAA | Bacteria | 0.0352 | 0.0150 | 0.0775 | 0.0010 | -0.9092 |
| 254 | GAA | Archaea | 0.0380 | 0.0165 | 0.0757 | 0.0048 | -0.7765 |
| 255 | GAA | Eukaryota | 0.0306 | 0.0111 | 0.0686 | 0.0030 | -0.9479 |
| 256 | GAG | All | 0.0283 | 0.0122 | 0.0869 | 0.0000 | 0.7692 |
| 257 | GAG | Bacteria | 0.0275 | 0.0117 | 0.0798 | 0.0000 | 0.7972 |
| 258 | GAG | Archaea | 0.0395 | 0.0174 | 0.0869 | 0.0068 | 0.8494 |
| 259 | GAG | Eukaryota | 0.0314 | 0.0097 | 0.0584 | 0.0055 | 0.8696 |
| 260 | GGA | All | 0.0152 | 0.0076 | 0.0460 | 0.0000 | -0.7375 |
| 261 | GGA | Bacteria | 0.0148 | 0.0079 | 0.0460 | 0.0000 | -0.7596 |
| 262 | GGA | Archaea | 0.0208 | 0.0082 | 0.0386 | 0.0053 | -0.8021 |
| 263 | GGA | Eukaryota | 0.0162 | 0.0040 | 0.0328 | 0.0029 | -0.3754 |
| 264 | GGC | All | 0.0283 | 0.0172 | 0.0734 | 0.0008 | 0.9331 |
| 265 | GGC | Bacteria | 0.0301 | 0.0177 | 0.0734 | 0.0008 | 0.9459 |
| 266 | GGC | Archaea | 0.0215 | 0.0134 | 0.0534 | 0.0015 | 0.9232 |
| 267 | GGC | Eukaryota | 0.0199 | 0.0103 | 0.0662 | 0.0009 | 0.9007 |
| 268 | GGG | All | 0.0123 | 0.0057 | 0.0513 | 0.0000 | 0.6654 |
| 269 | GGG | Bacteria | 0.0124 | 0.0056 | 0.0513 | 0.0000 | 0.6658 |
| 270 | GGG | Archaea | 0.0149 | 0.0069 | 0.0398 | 0.0018 | 0.7832 |
| 271 | GGG | Eukaryota | 0.0106 | 0.0045 | 0.0270 | 0.0007 | 0.6273 |
| 272 | GGT | All | 0.0158 | 0.0060 | 0.0443 | 0.0022 | -0.6421 |
| 273 | GGT | Bacteria | 0.0158 | 0.0062 | 0.0443 | 0.0022 | -0.6691 |
| 274 | GGT | Archaea | 0.0153 | 0.0060 | 0.0370 | 0.0030 | -0.5617 |
| 275 | GGT | Eukaryota | 0.0158 | 0.0044 | 0.0352 | 0.0038 | -0.3462 |
| 276 | TCA | All | 0.0091 | 0.0060 | 0.0422 | 0.0002 | -0.8642 |
| 277 | TCA | Bacteria | 0.0083 | 0.0056 | 0.0274 | 0.0002 | -0.9013 |
| 278 | TCA | Archaea | 0.0114 | 0.0065 | 0.0328 | 0.0008 | -0.8583 |
| 279 | TCA | Eukaryota | 0.0136 | 0.0054 | 0.0422 | 0.0021 | -0.8219 |
| 280 | TCC | All | 0.0108 | 0.0050 | 0.0331 | 0.0000 | 0.6298 |
| 281 | TCC | Bacteria | 0.0104 | 0.0049 | 0.0331 | 0.0000 | 0.6501 |
| 282 | TCC | Archaea | 0.0098 | 0.0040 | 0.0270 | 0.0021 | 0.6529 |
| 283 | TCC | Eukaryota | 0.0144 | 0.0041 | 0.0292 | 0.0018 | 0.7665 |
| 284 | TCG | All | 0.0108 | 0.0060 | 0.0455 | 0.0001 | 0.8124 |
| 285 | TCG | Bacteria | 0.0107 | 0.0059 | 0.0297 | 0.0004 | 0.8339 |
| 286 | TCG | Archaea | 0.0098 | 0.0068 | 0.0267 | 0.0001 | 0.8967 |
| 287 | TCG | Eukaryota | 0.0122 | 0.0059 | 0.0455 | 0.0007 | 0.6575 |
| 288 | TCT | All | 0.0094 | 0.0063 | 0.0367 | 0.0001 | -0.8528 |
| 289 | TCT | Bacteria | 0.0086 | 0.0061 | 0.0364 | 0.0001 | -0.9071 |
| 290 | TCT | Archaea | 0.0097 | 0.0057 | 0.0252 | 0.0006 | -0.8939 |
| 291 | TCT | Eukaryota | 0.0148 | 0.0048 | 0.0316 | 0.0023 | -0.6446 |
| 292 | TTC | All | 0.0189 | 0.0077 | 0.0398 | 0.0010 | 0.8507 |
| 293 | TTC | Bacteria | 0.0187 | 0.0079 | 0.0398 | 0.0010 | 0.8687 |
| 294 | TTC | Archaea | 0.0208 | 0.0073 | 0.0375 | 0.0048 | 0.8597 |
| 295 | TTC | Eukaryota | 0.0207 | 0.0045 | 0.0324 | 0.0051 | 0.7161 |
| 296 | TTT | All | 0.0219 | 0.0135 | 0.1003 | 0.0000 | -0.9387 |
| 297 | TTT | Bacteria | 0.0223 | 0.0140 | 0.1003 | 0.0000 | -0.9541 |
| 298 | TTT | Archaea | 0.0180 | 0.0114 | 0.0471 | 0.0009 | -0.9386 |
| 299 | TTT | Eukaryota | 0.0189 | 0.0082 | 0.0559 | 0.0041 | -0.9117 |
| 300 | TTA | All | 0.0153 | 0.0160 | 0.0851 | 0.0000 | -0.8746 |
| 301 | TTA | Bacteria | 0.0154 | 0.0162 | 0.0851 | 0.0000 | -0.8793 |
| 302 | TTA | Archaea | 0.0151 | 0.0141 | 0.0694 | 0.0001 | -0.8743 |
| 303 | TTA | Eukaryota | 0.0114 | 0.0111 | 0.0562 | 0.0002 | -0.9236 |
| 304 | TTG | All | 0.0153 | 0.0073 | 0.0528 | 0.0005 | -0.4940 |
| 305 | TTG | Bacteria | 0.0152 | 0.0075 | 0.0528 | 0.0005 | -0.5200 |
| 306 | TTG | Archaea | 0.0125 | 0.0059 | 0.0361 | 0.0027 | -0.5383 |
| 307 | TTG | Eukaryota | 0.0171 | 0.0058 | 0.0463 | 0.0033 | -0.4608 |
| 308 | TAC | All | 0.0132 | 0.0043 | 0.0374 | 0.0000 | 0.5661 |
| 309 | TAC | Bacteria | 0.0127 | 0.0040 | 0.0328 | 0.0000 | 0.6005 |
| 310 | TAC | Archaea | 0.0171 | 0.0061 | 0.0374 | 0.0045 | 0.7504 |
| 311 | TAC | Eukaryota | 0.0157 | 0.0036 | 0.0263 | 0.0038 | 0.6555 |
| 312 | TAT | All | 0.0167 | 0.0095 | 0.0601 | 0.0001 | -0.9271 |
| 313 | TAT | Bacteria | 0.0169 | 0.0096 | 0.0552 | 0.0001 | -0.9410 |
| 314 | TAT | Archaea | 0.0154 | 0.0095 | 0.0601 | 0.0008 | -0.9128 |
| 315 | TAT | Eukaryota | 0.0142 | 0.0077 | 0.0504 | 0.0010 | -0.9368 |
| 316 | TGC | All | 0.0066 | 0.0030 | 0.0245 | 0.0000 | 0.5759 |
| 317 | TGC | Bacteria | 0.0062 | 0.0027 | 0.0245 | 0.0000 | 0.6517 |
| 318 | TGC | Archaea | 0.0059 | 0.0027 | 0.0176 | 0.0008 | 0.3239 |
| 319 | TGC | Eukaryota | 0.0094 | 0.0032 | 0.0201 | 0.0014 | 0.5596 |
| 320 | TGT | All | 0.0048 | 0.0031 | 0.0247 | 0.0001 | -0.7338 |
| 321 | TGT | Bacteria | 0.0043 | 0.0027 | 0.0247 | 0.0001 | -0.8217 |
| 322 | TGT | Archaea | 0.0054 | 0.0026 | 0.0129 | 0.0006 | -0.6477 |
| 323 | TGT | Eukaryota | 0.0083 | 0.0037 | 0.0211 | 0.0009 | -0.7869 |
| 324 | TGG | All | 0.0121 | 0.0027 | 0.0280 | 0.0000 | 0.7476 |
| 325 | TGG | Bacteria | 0.0122 | 0.0026 | 0.0243 | 0.0000 | 0.7725 |
| 326 | TGG | Archaea | 0.0106 | 0.0018 | 0.0165 | 0.0055 | 0.4767 |
| 327 | TGG | Eukaryota | 0.0127 | 0.0025 | 0.0280 | 0.0046 | 0.6963 |
| 328 | ILE | All | 0.0639 | 0.0186 | 0.1767 | 0.0209 | -0.9183 |
| 329 | ILE | Bacteria | 0.0647 | 0.0184 | 0.1767 | 0.0209 | -0.9504 |
| 330 | ILE | Archaea | 0.0723 | 0.0205 | 0.1278 | 0.0257 | -0.9339 |
| 331 | ILE | Eukaryota | 0.0534 | 0.0126 | 0.1125 | 0.0229 | -0.9240 |
| 332 | LYS | All | 0.0576 | 0.0243 | 0.1894 | 0.0115 | -0.9264 |
| 333 | LYS | Bacteria | 0.0568 | 0.0248 | 0.1894 | 0.0115 | -0.9295 |
| 334 | LYS | Archaea | 0.0640 | 0.0271 | 0.1227 | 0.0129 | -0.8995 |
| 335 | LYS | Eukaryota | 0.0581 | 0.0148 | 0.1179 | 0.0268 | -0.8824 |
| 336 | ASN | All | 0.0381 | 0.0141 | 0.1303 | 0.0119 | -0.9194 |
| 337 | ASN | Bacteria | 0.0371 | 0.0134 | 0.1245 | 0.0119 | -0.9438 |
| 338 | ASN | Archaea | 0.0370 | 0.0128 | 0.0746 | 0.0168 | -0.9140 |
| 339 | ASN | Eukaryota | 0.0436 | 0.0155 | 0.1303 | 0.0198 | -0.8871 |
| 340 | TYR | All | 0.0301 | 0.0074 | 0.0650 | 0.0133 | -0.8672 |
| 341 | TYR | Bacteria | 0.0299 | 0.0075 | 0.0619 | 0.0155 | -0.8781 |
| 342 | TYR | Archaea | 0.0328 | 0.0062 | 0.0650 | 0.0225 | -0.6527 |
| 343 | TYR | Eukaryota | 0.0301 | 0.0058 | 0.0578 | 0.0133 | -0.8309 |
| 344 | PHE | All | 0.0410 | 0.0074 | 0.1061 | 0.0125 | -0.8306 |
| 345 | PHE | Bacteria | 0.0412 | 0.0076 | 0.1061 | 0.0125 | -0.8443 |
| 346 | PHE | Archaea | 0.0392 | 0.0058 | 0.0573 | 0.0226 | -0.7546 |
| 347 | PHE | Eukaryota | 0.0397 | 0.0053 | 0.0648 | 0.0183 | -0.8018 |
| 348 | MET | All | 0.0251 | 0.0037 | 0.0425 | 0.0135 | -0.2492 |
| 349 | MET | Bacteria | 0.0253 | 0.0037 | 0.0375 | 0.0135 | -0.2835 |
| 350 | MET | Archaea | 0.0251 | 0.0041 | 0.0377 | 0.0161 | -0.3884 |
| 351 | MET | Eukaryota | 0.0236 | 0.0025 | 0.0425 | 0.0148 | -0.1917 |
| 352 | LEU | All | 0.0990 | 0.0075 | 0.1473 | 0.0733 | 0.3244 |
| 353 | LEU | Bacteria | 0.1002 | 0.0071 | 0.1473 | 0.0781 | 0.3659 |
| 354 | LEU | Archaea | 0.0946 | 0.0077 | 0.1171 | 0.0733 | 0.0808 |
| 355 | LEU | Eukaryota | 0.0916 | 0.0053 | 0.1390 | 0.0767 | -0.0629 |
| 356 | SER | All | 0.0614 | 0.0096 | 0.1064 | 0.0338 | -0.4210 |
| 357 | SER | Bacteria | 0.0586 | 0.0063 | 0.0921 | 0.0338 | -0.6545 |
| 358 | SER | Archaea | 0.0610 | 0.0070 | 0.0805 | 0.0442 | -0.3691 |
| 359 | SER | Eukaryota | 0.0812 | 0.0061 | 0.1064 | 0.0568 | -0.0079 |
| 360 | GLN | All | 0.0348 | 0.0082 | 0.1804 | 0.0113 | -0.0925 |
| 361 | GLN | Bacteria | 0.0346 | 0.0075 | 0.0704 | 0.0113 | -0.1348 |
| 362 | GLN | Archaea | 0.0244 | 0.0050 | 0.0420 | 0.0121 | 0.0050 |
| 363 | GLN | Eukaryota | 0.0403 | 0.0097 | 0.1804 | 0.0209 | 0.0844 |
| 364 | GLU | All | 0.0638 | 0.0085 | 0.1174 | 0.0281 | -0.4623 |
| 365 | GLU | Bacteria | 0.0633 | 0.0080 | 0.0967 | 0.0281 | -0.5361 |
| 366 | GLU | Archaea | 0.0785 | 0.0121 | 0.1174 | 0.0554 | 0.1546 |
| 367 | GLU | Eukaryota | 0.0622 | 0.0053 | 0.0963 | 0.0388 | -0.3936 |
| 368 | CYS | All | 0.0113 | 0.0040 | 0.0333 | 0.0016 | -0.1393 |
| 369 | CYS | Bacteria | 0.0104 | 0.0029 | 0.0333 | 0.0016 | -0.1450 |
| 370 | CYS | Archaea | 0.0112 | 0.0027 | 0.0189 | 0.0061 | -0.2769 |
| 371 | CYS | Eukaryota | 0.0178 | 0.0046 | 0.0320 | 0.0094 | -0.2465 |
| 372 | THR | All | 0.0533 | 0.0054 | 0.0768 | 0.0207 | 0.2728 |
| 373 | THR | Bacteria | 0.0529 | 0.0051 | 0.0730 | 0.0207 | 0.2720 |
| 374 | THR | Archaea | 0.0514 | 0.0082 | 0.0768 | 0.0353 | 0.5098 |
| 375 | THR | Eukaryota | 0.0566 | 0.0050 | 0.0699 | 0.0375 | 0.3193 |
| 376 | ASP | All | 0.0535 | 0.0059 | 0.0892 | 0.0216 | 0.3904 |
| 377 | ASP | Bacteria | 0.0533 | 0.0055 | 0.0888 | 0.0216 | 0.4380 |
| 378 | ASP | Archaea | 0.0574 | 0.0132 | 0.0892 | 0.0360 | 0.5227 |
| 379 | ASP | Eukaryota | 0.0532 | 0.0044 | 0.0758 | 0.0319 | -0.0011 |
| 380 | HIS | All | 0.0206 | 0.0035 | 0.0357 | 0.0097 | 0.4462 |
| 381 | HIS | Bacteria | 0.0203 | 0.0032 | 0.0346 | 0.0104 | 0.5062 |
| 382 | HIS | Archaea | 0.0176 | 0.0024 | 0.0243 | 0.0097 | 0.5722 |
| 383 | HIS | Eukaryota | 0.0246 | 0.0025 | 0.0357 | 0.0144 | 0.3402 |
| 384 | VAL | All | 0.0705 | 0.0094 | 0.1103 | 0.0209 | 0.7249 |
| 385 | VAL | Bacteria | 0.0715 | 0.0088 | 0.1103 | 0.0209 | 0.7698 |
| 386 | VAL | Archaea | 0.0763 | 0.0117 | 0.1095 | 0.0512 | 0.8079 |
| 387 | VAL | Eukaryota | 0.0618 | 0.0057 | 0.0790 | 0.0386 | 0.6323 |
| 388 | TRP | All | 0.0122 | 0.0026 | 0.0282 | 0.0000 | 0.7417 |
| 389 | TRP | Bacteria | 0.0122 | 0.0026 | 0.0244 | 0.0000 | 0.7726 |
| 390 | TRP | Archaea | 0.0105 | 0.0017 | 0.0160 | 0.0056 | 0.5031 |
| 391 | TRP | Eukaryota | 0.0128 | 0.0025 | 0.0282 | 0.0046 | 0.6940 |
| 392 | ARG | All | 0.0574 | 0.0153 | 0.1212 | 0.0202 | 0.9362 |
| 393 | ARG | Bacteria | 0.0576 | 0.0159 | 0.1212 | 0.0202 | 0.9464 |
| 394 | ARG | Archaea | 0.0573 | 0.0135 | 0.0956 | 0.0298 | 0.8499 |
| 395 | ARG | Eukaryota | 0.0577 | 0.0090 | 0.0854 | 0.0275 | 0.8247 |
| 396 | GLY | All | 0.0721 | 0.0114 | 0.1046 | 0.0224 | 0.9025 |
| 397 | GLY | Bacteria | 0.0735 | 0.0108 | 0.1046 | 0.0224 | 0.9446 |
| 398 | GLY | Archaea | 0.0729 | 0.0084 | 0.0934 | 0.0500 | 0.9150 |
| 399 | GLY | Eukaryota | 0.0627 | 0.0103 | 0.1028 | 0.0301 | 0.8792 |
| 400 | PRO | All | 0.0444 | 0.0100 | 0.0807 | 0.0132 | 0.8532 |
| 401 | PRO | Bacteria | 0.0435 | 0.0095 | 0.0704 | 0.0132 | 0.9276 |
| 402 | PRO | Archaea | 0.0409 | 0.0058 | 0.0543 | 0.0279 | 0.8499 |
| 403 | PRO | Eukaryota | 0.0530 | 0.0098 | 0.0807 | 0.0206 | 0.8471 |
| 404 | ALA | All | 0.0898 | 0.0274 | 0.1653 | 0.0111 | 0.9389 |
| 405 | ALA | Bacteria | 0.0928 | 0.0275 | 0.1653 | 0.0111 | 0.9561 |
| 406 | ALA | Archaea | 0.0756 | 0.0205 | 0.1251 | 0.0362 | 0.9359 |
| 407 | ALA | Eukaryota | 0.0757 | 0.0204 | 0.1621 | 0.0222 | 0.9013 |
| 408 | K+R | All | 0.1150 | 0.0133 | 0.2123 | 0.0793 | -0.6147 |
| 409 | K+R | Bacteria | 0.1144 | 0.0133 | 0.2123 | 0.0805 | -0.6043 |
| 410 | K+R | Archaea | 0.1213 | 0.0197 | 0.1632 | 0.0793 | -0.6568 |
| 411 | K+R | Eukaryota | 0.1158 | 0.0087 | 0.1549 | 0.0869 | -0.6426 |
| 412 | K+R+H | All | 0.1357 | 0.0118 | 0.2227 | 0.0992 | -0.5657 |
| 413 | K+R+H | Bacteria | 0.1347 | 0.0115 | 0.2227 | 0.1003 | -0.5581 |
| 414 | K+R+H | Archaea | 0.1389 | 0.0182 | 0.1767 | 0.0992 | -0.6310 |
| 415 | K+R+H | Eukaryota | 0.1404 | 0.0084 | 0.1754 | 0.1102 | -0.5684 |
| 416 | D+E | All | 0.1173 | 0.0099 | 0.1861 | 0.0534 | -0.1645 |
| 417 | D+E | Bacteria | 0.1167 | 0.0085 | 0.1707 | 0.0534 | -0.2243 |
| 418 | D+E | Archaea | 0.1359 | 0.0207 | 0.1861 | 0.1037 | 0.4238 |
| 419 | D+E | Eukaryota | 0.1154 | 0.0067 | 0.1424 | 0.0730 | -0.3170 |
| 420 | F+L+I+V+M | All | 0.2995 | 0.0205 | 0.3949 | 0.1756 | -0.7249 |
| 421 | F+L+I+V+M | Bacteria | 0.3030 | 0.0173 | 0.3949 | 0.2497 | -0.9022 |
| 422 | F+L+I+V+M | Archaea | 0.3075 | 0.0215 | 0.3496 | 0.2602 | -0.6987 |
| 423 | F+L+I+V+M | Eukaryota | 0.2701 | 0.0171 | 0.3475 | 0.1756 | -0.7651 |
| 424 | hydrophobics | All | 0.4015 | 0.0222 | 0.4625 | 0.3080 | 0.5749 |
| 425 | hydrophobics | Bacteria | 0.4080 | 0.0157 | 0.4625 | 0.3408 | 0.8106 |
| 426 | hydrophobics | Archaea | 0.3936 | 0.0170 | 0.4359 | 0.3302 | 0.2934 |
| 427 | hydrophobics | Eukaryota | 0.3586 | 0.0123 | 0.4255 | 0.3080 | 0.5664 |
| 428 | Ile (ATA+ATT) | All | 0.0377 | 0.0263 | 0.1746 | 0.0001 | -0.9627 |
| 429 | Ile (ATA+ATT) | Bacteria | 0.0375 | 0.0268 | 0.1746 | 0.0001 | -0.9708 |
| 430 | Ile (ATA+ATT) | Archaea | 0.0484 | 0.0282 | 0.1234 | 0.0024 | -0.9609 |
| 431 | Ile (ATA+ATT) | Eukaryota | 0.0320 | 0.0166 | 0.1035 | 0.0027 | -0.9638 |
| 432 | Ile (ATC) | All | 0.0258 | 0.0102 | 0.0619 | 0.0011 | 0.8142 |
| 433 | Ile (ATC) | Bacteria | 0.0268 | 0.0104 | 0.0619 | 0.0011 | 0.8218 |
| 434 | Ile (ATC) | Archaea | 0.0232 | 0.0106 | 0.0542 | 0.0037 | 0.7551 |
| 435 | Ile (ATC) | Eukaryota | 0.0212 | 0.0058 | 0.0352 | 0.0042 | 0.7561 |
| 436 | Ser (TCn) | All | 0.0401 | 0.0080 | 0.0818 | 0.0139 | -0.3136 |
| 437 | Ser (TCn) | Bacteria | 0.0380 | 0.0057 | 0.0655 | 0.0139 | -0.4314 |
| 438 | Ser (TCn) | Archaea | 0.0407 | 0.0066 | 0.0573 | 0.0236 | -0.3060 |
| 439 | Ser (TCn) | Eukaryota | 0.0549 | 0.0069 | 0.0818 | 0.0315 | -0.0651 |
| 440 | Ser (AGT+AGC) | All | 0.0212 | 0.0043 | 0.0574 | 0.0074 | -0.3256 |
| 441 | Ser (AGT+AGC) | Bacteria | 0.0206 | 0.0040 | 0.0420 | 0.0074 | -0.3879 |
| 442 | Ser (AGT+AGC) | Archaea | 0.0203 | 0.0033 | 0.0312 | 0.0117 | -0.1318 |
| 443 | Ser (AGT+AGC) | Eukaryota | 0.0260 | 0.0038 | 0.0574 | 0.0153 | 0.1340 |
| 444 | Arg (CGn) | All | 0.0432 | 0.0223 | 0.1047 | 0.0002 | 0.9354 |
| 445 | Arg (CGn) | Bacteria | 0.0452 | 0.0227 | 0.1047 | 0.0007 | 0.9515 |
| 446 | Arg (CGn) | Archaea | 0.0277 | 0.0217 | 0.0886 | 0.0002 | 0.8659 |
| 447 | Arg (CGn) | Eukaryota | 0.0367 | 0.0135 | 0.0774 | 0.0016 | 0.8265 |
| 448 | Arg (AGA+AGG) | All | 0.0144 | 0.0102 | 0.0694 | 0.0009 | -0.6290 |
| 449 | Arg (AGA+AGG) | Bacteria | 0.0127 | 0.0091 | 0.0621 | 0.0009 | -0.7029 |
| 450 | Arg (AGA+AGG) | Archaea | 0.0300 | 0.0162 | 0.0694 | 0.0023 | -0.4403 |
| 451 | Arg (AGA+AGG) | Eukaryota | 0.0209 | 0.0071 | 0.0543 | 0.0017 | -0.5186 |

## References

1. Agashe D, Shankar N. The evolution of bacterial DNA base composition. J Exp Zool B Mol Dev Evol. 2014 Nov;322(7):517–528.

2. Marin A, Xia X. GC skew in protein-coding genes between the leading and lagging strands in bacterial genomes: new substitution models incorporating strand bias. J Theor Biol. 2008 Aug;253(3):508–513.

3. Hildebrand F, Meyer A, Eyre-Walker A. Evidence of selection upon genomic GC-content in bacteria. PLoS Genet. 2010 Sep;6(9):e1001107.

4. Reichenberger ER, Rosen G, Hershberg U, Hershberg R. Prokaryotic nucleotide composition is shaped by both phylogeny and the environment. Genome Biol Evol. 2015 Apr;7(5):1380–1389.

5. Kennedy SP, Ng WV, Salzberg SL, Hood L, DasSarma S. Understanding the adaptation of Halobacterium species NRC-1 to its extreme environment through computational analysis of its genome sequence. Genome Res. 2001 Oct;11(10):1641–1650.

6. Prat Y, Fromer M, Linial N, Linial M. Codon usage is associated with the evolutionary age of genes in metazoan genomes. BMC Evol Biol. 2009;9:285.

7. Wu H, Zhang Z, Hu S, Yu J. On the molecular mechanism of GC content variation among eubacterial genomes. Biol Direct. 2012 Jan;7:2.

8. Knight RD, Freeland SJ, Landweber LF. A simple model based on mutation and selection explains trends in codon and amino-acid usage and GC composition within and across genomes. Genome Biol. 2001;2(4):RESEARCH0010.

9. Peng Z, Uversky VN, Kurgan L. Genes encoding intrinsic disorder in Eukaryota have high GC content. Intrinsically Disord Proteins. 2016;4(1):e1262225.

10. Du MZ, Zhang C, Wang H, Liu S, Wei W, Guo FB. The GC Content as a Main Factor Shaping the Amino Acid Usage During Bacterial Evolution Process. Front Microbiol. 2018;9:2948.

11. Basile W, Sachenkova O, Light S, Elofsson A. High GC content causes orphan proteins to be intrinsically disordered. PLOS Computational Biology. 2017 03;13(3):1–19. Available from: https://doi.org/10.1371/journal.pcbi.1005375.

12. Basile W, Sachenkova O, Light S, Elofsson A. High GC content causes orphan proteins to be intrinsically disordered. PLoS Comput Biol. 2017 Mar;13(3):e1005375.

13. Mangiapane E, Pessione A, Pessione E. Selenium and selenoproteins: an overview on different biological systems. Curr Protein Pept Sci. 2014;15(6):598–607.

14. Wang YN, Ji WH, Li XR, Liu YS, Zhou JH. Unique features of nucleotide and codon usage patterns in mycoplasmas revealed by information entropy. Biosystems. 2018 Mar;165:1–7.

15. Saier MH Jr. Understanding the Genetic Code. J Bacteriol. 2019 Aug;201(15).

16. Singer GA, Hickey DA. Nucleotide bias causes a genomewide bias in the amino acid composition of proteins. Mol Biol Evol. 2000 Nov;17(11):1581–1588.

17. Pandya S, Struck TJ, Mannakee BK, Paniscus M, Gutenkunst RN. Testing whether metazoan tyrosine loss was driven by selection against promiscuous phosphorylation. Mol Biol Evol. 2015 Jan;32(1):144–152.

18. Tekaia F, Yeramian E. Evolution of proteomes: fundamental signatures and global trends in amino acid compositions. BMC Genomics. 2006 Dec;7:307.

19. Akashi H, Gojobori T. Metabolic efficiency and amino acid composition in the proteomes of Escherichia coli and Bacillus subtilis. Proc Natl A cad Sci U S A. 2002 Mar;99(6):3695–3700.

20. Raiford DW, Heizer EM Jr, Miller RV, Akashi H, Raymer ML, Krane DE. Do amino acid biosynthetic costs constrain protein evolution in Saccharomyces cerevisiae? J Mol Evol. 2008 Dec;67(6):621–630.

21. de Lorenzo V, Sekowska A, Danchin A. Chemical reactivity drives spatiotemporal organisation of bacterial metabolism. FEMS Microbiol Rev. 2015 Jan;39(1):96–119.

22. Goldstein RA. Population size dependence of fitness effect distribution and substitution rate probed by biophysical model of protein thermostability. Genome Biol Evol. 2013;5(9):1584–1593.

23. Consortium TU. The Universal Protein Resource (UniProt) in 2010. Nucleic Acids Res. 2010 Jan;38(Database issue):D142–8. Available from: http://view.ncbi.nlm.nih.gov/pubmed/19843607.

24. Elofsson A. Dataset for paper. 2018 12;Available from: https://figshare.com/articles/Dataset_for_paper/7478381.

25. Muller KE, Fetterman BA. Regression and ANOVA: An Integrated Approach Using SAS Software. New York, NY, USA: John Wiley & Sons, Inc.; 2003.

26. In: Kirch W, editor. Pearson’s Correlation Coefficient. Dordrecht: Springer Netherlands; 2008. p. 1090–1091. Available from: https://doi.org/10.1007/978-1-4020-5614-7_2569.

27. Pedregosa F, Varoquaux G, Gramfort A, Michel V, Thirion B, Grisel O, et al. Scikit-learn: Machine Learning in Python. Journal of Machine Learning Research. 2011;12:2825–2830.

28. Clement Y, Fustier MA, Nabholz B, Glemin S. The bimodal distribution of genic GC content is ancestral to monocot species. Genome Biol Evol. 2015 Jan;7(1):336–348.

